# Host-to-symbiont transfer of fatty-acid-derived carbon in the *Paramecium bursaria*-*Chlorella variabilis* photosymbiosis

**DOI:** 10.1101/2025.11.27.690892

**Authors:** Yan-Jun Chen, Chi-Yen Wei, Chia-Wei Hsu, Pei-Ling Wang, Silver Sung-Yun Hsiao, Der-Chuen Lee, Chia-Ling Yang, Chuan Ku, Md Mostafa Kamal, Jun-Yi Leu

**Affiliations:** Institute of Molecular Biology, Academia Sinica, Taipei, Taiwan; Agricultural Biotechnology Research Center, Academia Sinica, Taipei, Taiwan; Institute of Oceanography, National Taiwan University, Taipei, Taiwan; Institute of Astronomy and Astrophysics, Academia Sinica, Taipei, Taiwan; Institute of Earth Sciences, Academia Sinica, Taipei, Taiwan; Institute of Plant and Microbial Biology, Academia Sinica, Taipei, Taiwan

**Keywords:** endosymbiosis, transcriptome, lipidome, fatty-acid carbon transfer, ciliate

## Abstract

Metabolic exchange stabilizes photosymbioses, but carbon transfer is usually framed as exporting photosynthate from algal symbionts to heterotrophic hosts. Previous work showed that *Paramecium bursaria* reorganizes its lipid droplets during symbiosis and that perturbing host lipid metabolism reduces endosymbiont abundance. Here, we investigated reciprocal organic-carbon exchange between *P. bursaria* and its intracellular *Chlorella* symbionts. Symbiosis was associated with extensive lipid remodeling in both partners and marked triacylglycerol accumulation in endosymbiotic algae. Following host supplementation with ^13^C-oleic acid, NanoSIMS and EA-IRMS detected ^13^C enrichment in intracellular algae. Isotopologue-resolved lipidomics further showed incorporation of host-acquired fatty-acid carbon into algal storage and membrane lipids. These findings reveal an unexpected host-to-symbiont direction of organic-carbon transfer and expand the conventional model of metabolic exchange in photosymbiosis.

## Introduction

Symbiotic associations enable distinct species to integrate complementary metabolic capacities, thereby expanding their ecological potential in diverse environments^1,2^. Among these interactions, photosymbioses are particularly widespread and ecologically important, occurring in ciliates, corals, hydras, jellyfish and sea anemones^3^. In these systems, symbiosis is commonly viewed as a nutritional partnership in which photosynthetic symbionts provide fixed carbon to heterotrophic hosts, whereas hosts supply inorganic nutrients and a protected intracellular environment^4–8^. However, the extent to which carbon exchange is reciprocal, and how organic metabolites move across the host-symbiont interface, remain incompletely understood. Defining the directionality, molecular form, and cellular route of metabolite exchange is therefore essential for understanding how photosymbiotic associations are stabilized.

Lipids are increasingly recognized as active mediators of symbiotic physiology rather than passive storage molecules^9,10^. In cnidarian-algal symbioses, for example, lipid abundance and peroxidation levels closely reflect symbiotic state^10^, and sterol transfer from algae to the host is essential because cnidarian hosts are sterol auxotrophs^11^. In plant-fungal mutualisms, host-derived lipids are transferred to fungal partners, contributing to partner dependency and long-term persistence^12–15^. These observations suggest that lipids may represent important currencies of metabolic integration across evolutionarily diverse symbioses. However, in most photosymbiotic systems, carbon-rich organic metabolites are still primarily considered to move from photosymbionts to their partners. Whether non-photosynthetic partners can also provide organic carbon to support or regulate photosymbiont metabolism remains unresolved.

Ciliates are an ancient and widespread group of protists found in various environments. Many ciliates form endosymbioses with algae or bacteria, which allow them to explore novel ecological niches^16,17^. Among them, *Paramecium bursaria* and its intracellular green alga *Chlorella* provide a tractable model for investigating host-symbiont nutrient trafficking^18,19^. Both partners can be maintained independently in culture, and aposymbiotic *P. bursaria* cells can be experimentally reinfected with isolated algae^20^. Previous studies have shown that the host supplies nitrogenous compounds that support algal metabolism, whereas the alga fixes inorganic carbon and transfers photosynthates to the host^21–23^. More recently, this association has been proposed to operate as an integrated nutrient-recycling system that promotes survival under nutrient-poor conditions^24^. In our previous study, we found that lipid droplets are reorganized in symbiotic *P. bursaria* cells and that pharmacological perturbation of lipid metabolism in the host cell significantly reduces the endosymbiont number^25^. These observations suggest that host lipid metabolism contributes directly to the maintenance of intracellular algae. However, it remains unaddressed whether the host cell and endosymbiotic algae have exchanged lipids or fatty acids and how these metabolic processes are integrated.

Recent advances in multi-omics approaches provide powerful tools for investigating bidirectional nutrient exchange in symbiotic systems. By integrating transcriptomics and metabolomics, it becomes possible to simultaneously track host- and symbiont-derived pathways, identify candidate metabolites involved in reciprocal exchange, and resolve their regulation under different environmental conditions^26–28^. In this study, we combined comparative transcriptomics, lipidomics, imaging mass spectrometry and stable-isotope tracing to investigate how lipid metabolism contributes to the establishment of endosymbiosis between *P. bursaria* and *Chlorella*. We found that symbiosis is associated with extensive remodeling of algal lipid metabolic pathways and enrichment of multiple lipid classes, including triacylglycerols. We further detected incorporation of host-derived fatty-acid carbon into lipids of endosymbiotic algae, supporting a model in which host-to-symbiont carbon transfer contributes to metabolic reciprocity in the *P. bursaria*-*Chlorella* association. Together, these findings expand the conventional view of nutrient exchange in photosymbiosis and identify host-derived fatty acids as a potential contributor to symbiotic stability.

## Result

### Characterization of lipid droplet components in *Paramecium bursaria*

Our previous study revealed that the morphology and protein composition of lipid droplets differ significantly between symbiotic and aposymbiotic *P. bursaria* cells. Moreover, the triacylglycerol (TG) biosynthetic pathway is upregulated in symbiotic host cells^25^.

To characterize the composition of host lipid droplets in detail, we isolated lipid droplet-enriched fractions from both symbiotic and aposymbiotic *P. bursaria* cells and performed comprehensive lipidomic analysis using liquid chromatography-tandem mass spectrometry (LC-MS/MS) (see Materials and Methods for details). We analyzed four biological replicates for each sample. Within-condition Pearson correlations were consistently high (r = 0.90-0.97), supporting replicate consistency (Supporting Fig. S1a; Supporting Data 1, A). Between-condition correlations were lower (r = 0.76-0.83), indicating a condition-associated shift in host lipid composition (Supporting Fig. S1a). Furthermore, principal-component analysis (PCA) provided a complementary multivariate view and separated symbiotic from aposymbiotic samples along PC1 (74.3% of variance; exact-label PERMANOVA pseudo-F = 15.77, *P* = 0.042; n = 4 biological replicates per group; Fig. 1a). Consistent with this observation, the total lipid abundance in symbiotic *P. bursaria* lipid droplets was approximately 8.15-fold higher than that in aposymbiotic cells (two-sided Welch test, *P* = 0.0269; Fig. 1b; Supporting Table S1a). Analysis of lipid class distribution also showed that TG was the dominant lipid subclass in both conditions, accounting for 91.1% and 59.3% of total lipid content in symbiotic and aposymbiotic cells, respectively (Fig. 1c; Supporting Data 1, B; Supporting Table S1b). In total, 453 host lipid species across 18 subclasses had nonzero mean abundance in both conditions and were included in the fold-change analysis (Supporting Fig. S1b,c; Supporting Data 1, C; Supporting Table S1c). This comparison set contained 214 triacylglycerol species, representing the largest proportion of the detected lipidome. Other major subclasses included phosphatidylethanolamine (PE), phosphatidylcholine (PC), ether-linked TG (etherTG), and ether-linked PC (etherPC), each consisting of approximately 20-40 lipid species (Supporting Fig. S1b).

**Figure 1.**
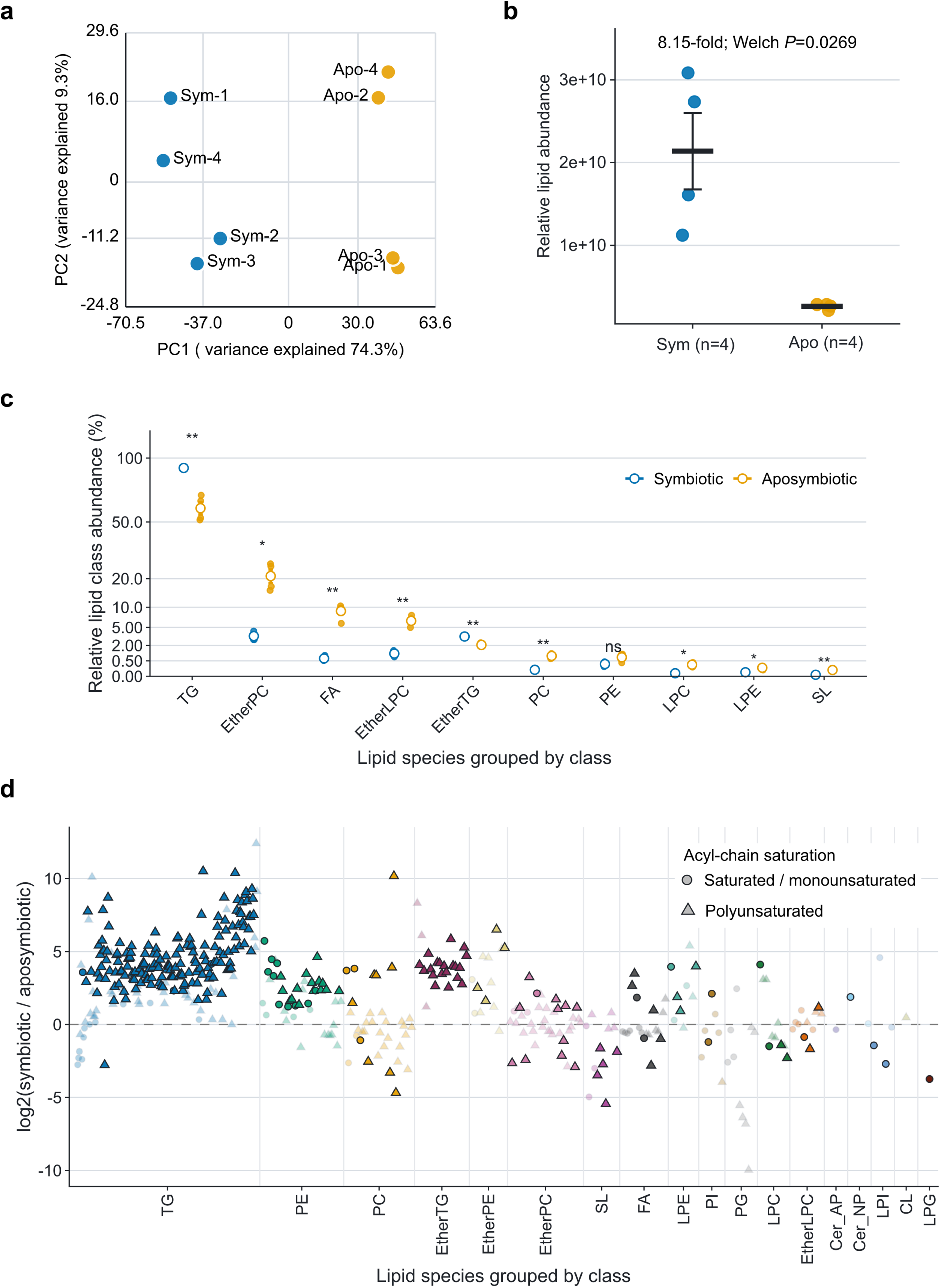
Symbiosis is associated with lipidome-wide remodeling of host lipid droplet-enriched fractions. **(a)** PCA of log_2_-transformed lipid-species abundance. Points are biological replicates (n = 4 per state); the annotation reports exact-label PERMANOVA pseudo-F = 15.77 and *P* = 0.042. **(b)** Total lipid abundance. Points are biological replicates; crossbars and error bars show mean ± SEM (8.15-fold; two-sided Welch test, *P* = 0.0269). **(c)** Relative abundance of the ten lipid classes with a mean abundance of at least 0.05% in either state. Points are biological replicates, open circles show means and error bars show SEM; the y axis is square-root scaled. Asterisks denote BH-adjusted two-sided Welch tests (\**P* < 0.05, \*\**P* < 0.01, \*\*\**P* < 0.001 and \*\*\*\**P* < 0.0001; ns, *P* ≥ 0.05). **(d)** Species-level symbiotic/aposymbiotic log_2_ fold changes for the 453 annotations eligible for fold-change analysis, arranged by lipid class. Fill color denotes lipid class, shape denotes acyl-chain saturation category, and opacity/outline denotes nominal two-sided Welch *P* < 0.05 versus *P* ≥ 0.05; marker size is constant. The horizontal dashed line marks no change, and horizontal positions are categorical and compressed within each class. Supporting Data 1, B-C; Supporting Tables S1a-S1d.

We next examined how changes in host lipid composition were distributed across lipid classes and acyl chain features. In symbiotic host lipid droplet-enriched fractions, triacylglycerol abundance increased by approximately 12.7-fold (two-sided Welch test on log_2_-transformed class-summed abundance, *P* = 0.00123, BH-adjusted *P* = 0.00607; Supporting Fig. S1c; Supporting Table S1d). Many TG species with positive fold changes contained polyunsaturated fatty-acid (PUFA) chains (Fig. 1d; Supporting Data 1, C). Consistently, subclass-level comparisons showed that most TG species were markedly more abundant in symbiotic *P. bursaria* (Fig. 1d; Supporting Data 1, C; Supporting Table S1d), indicating extensive remodeling of neutral lipid storage.

The lipid-class summary confirmed that host remodeling extended beyond TG to ether lipids, phospholipids, and sphingolipids, rather than representing a uniform increase across classes (Supporting Fig. S1c; Supporting Data 1, C; Supporting Table S1d). Ether-linked neutral lipids, including etherTG and etherPE, were also enriched, suggesting that symbiosis remodels not only neutral lipid storage but also lipid subclasses often associated with membrane reorganization and stress-related lipid metabolism^29,30^. By contrast, phospholipid changes were selective rather than uniform: PE increased, whereas PG and lysophosphatidylglycerol (LPG) decreased. Sphingolipid-related species also showed varying patterns in which sulfonolipids (SL) decreased and ceramide-NP increased. Together, these patterns show selective differences in the host lipid-droplet lipidome between symbiotic and aposymbiotic cells, spanning TG, ether lipids, phospholipids and sphingolipid subclasses.

### Host-acquired fatty-acid carbon reaches endosymbiotic algae

The spatial proximity between host lipid droplets and endosymbiotic algae in a scanning electron microscopy (SEM) image of a thin section supports a physical context in which host lipid pools could contribute to interpartner exchange (Supporting Fig. S2a)^25^. To test whether host-acquired lipid carbon can reach the endosymbiont, we designed an experimental workflow in which symbiotic *P. bursaria* cells were supplemented with oleic acid with or without ^13^C-labeling for 24 hours, followed by imaging- and isotope-based analyses of both host and algal fractions (Fig. 2a).

**Figure 2.**
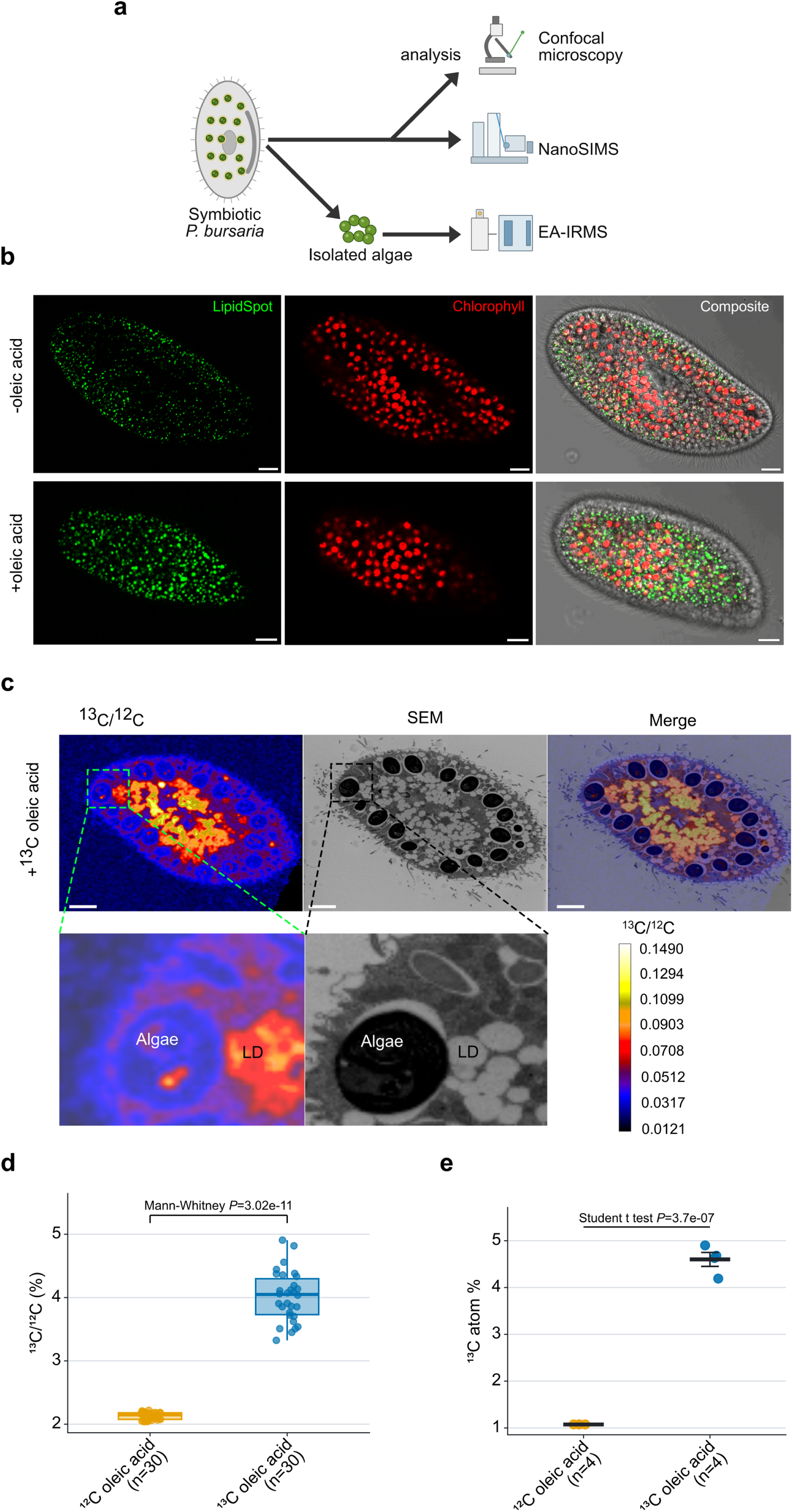
Experimental workflow and evidence for host-to-symbiont fatty-acid-carbon transfer. **(a)** Oleic-acid feeding and isotope-tracing workflow. **(b)** Representative confocal images without oleic acid (upper row) and after oleic-acid supplementation (lower row). LipidSpot signal is green and chlorophyll autofluorescence is red; scale bars, 10 µM. **(c)** Representative NanoSIMS ^13^C/^12^C maps, corresponding SEM images of thin sections and merged images after host feeding with ^13^C_18_-oleic acid. Lower panels show the boxed region at higher magnification; scale bars, 5 µM. **(d)** Algal ^13^C/^12^C ratios. Points are individual analyzed cells treated as experimental units (n = 30 per group); boxes show the median and interquartile range, and whiskers extend to 1.5 times the interquartile range (two-sided Mann-Whitney U test, *P* = 3.02 × 10^-11^). **(e)** EA-IRMS ^13^C atom percentage. Points are biological replicates (n = 4 per group); crossbars and error bars show mean ± SEM (two-sided equal-variance Student t test, *P* = 3.70 × 10^-7^). Supporting Data 2, A; Supporting Tables S2a-S2b.

We first asked whether exogenous fatty acids can be incorporated into host lipid stores. After oleic acid supplementation, symbiotic *P. bursaria* cells displayed enlarged lipid droplets relative to untreated controls (Fig. 2b), consistent with incorporation of the supplied fatty acid into host neutral-lipid pools.

We next traced the fate of host-supplied fatty acids using nanoscale secondary ion mass spectrometry (NanoSIMS). NanoSIMS provided nanoscale maps of isotopic signals within *P. bursaria* cells. NanoSIMS imaging revealed strong ^13^C enrichment within host lipid droplets and a concomitant increase in the ^13^C/^12^C ratio in the intracellular *Chlorella* cells relative to the unlabeled control (two-sided Mann-Whitney U test, *P* = 3.02 × 10^-11^; n = 30 analyzed cells per group; Fig. 2c and 2d; Supporting Data 2, A; Supporting Table S2a). This spatial enrichment shows that carbon derived from host-acquired oleate reached intracellular algae under the assay conditions.

We used an elemental analyzer coupled with isotope ratio mass spectrometry (EA-IRMS) to further determine the exact isotope composition of endosymbiotic algae isolated from *P. bursaria* cells. Quantitative elemental analysis confirmed significantly elevated ^13^C atom percentages in algae isolated from symbiotic *P. bursaria* fed with ^13^C-labeled oleic acid (two-sided equal-variance Student t-test, *P* = 3.70 × 10^-7^; n = 4 biological replicates per group; Fig. 2e; Supporting Table S2b). Together, the imaging and bulk isotope measurements show that carbon derived from host-acquired oleate reached intracellular algal endosymbionts, although the molecular form that crossed the host-symbiont interface remains unresolved.

### Stable-isotope tracing reveals incorporation of host-derived fatty-acid carbon into symbiont lipids

Having established direct host-to-symbiont carbon transfer, we next asked which algal lipid species incorporate host-derived carbon compounds. To address this question, we supplied *P. bursaria* with ^13^C-labeled oleic acid and subsequently extracted lipids from endosymbiotic *C. variabilis* cells for LC-MS/MS analysis (see Materials and Methods for details). In brief, lipidomic profiling was performed in both positive and negative ionization modes to ensure broad lipid coverage. Peak annotation for unlabeled samples was conducted using MS-DIAL^31^, whereas isotopologue extraction and labeling fraction quantification were performed using El-Maven^32^. Isotopic natural abundance correction was applied using IsoCorrectoR^33^.

Following host supplementation with ^13^C_18_-oleic acid, 189 of 772 structural lipid annotations in endosymbiotic *C. variabilis* met the prespecified labeling criteria (delta labeling > 0.05 and isotope-labeling BH-adjusted *P* < 0.05; Fig. 3a; Supporting Data 2, B; Supporting Table S3a). After duplicate structural annotations were collapsed to 631 unique base/sum-composition annotations, 165 remained labeled; class-versus-rest Fisher tests identified TG as overrepresented (BH-adjusted *P* = 0.000323) and ceramide as underrepresented (BH-adjusted *P* = 0.0340) among labeled annotations (Fig. 3b; Supporting Data 2, C; Supporting Table S3b). The corresponding structural-entry analysis gave the same directions for TG (BH-adjusted *P* = 0.00580) and ceramide (BH-adjusted *P* = 0.0149), showing that these class-level conclusions were retained before duplicate annotations were collapsed (Supporting Fig. S2b). The fraction of labeled annotations varied among lipid classes, indicating that oleate-derived carbon was incorporated unevenly across the algal lipidome (Fig. 3a and Supporting Fig. S2c).

**Figure 3.**
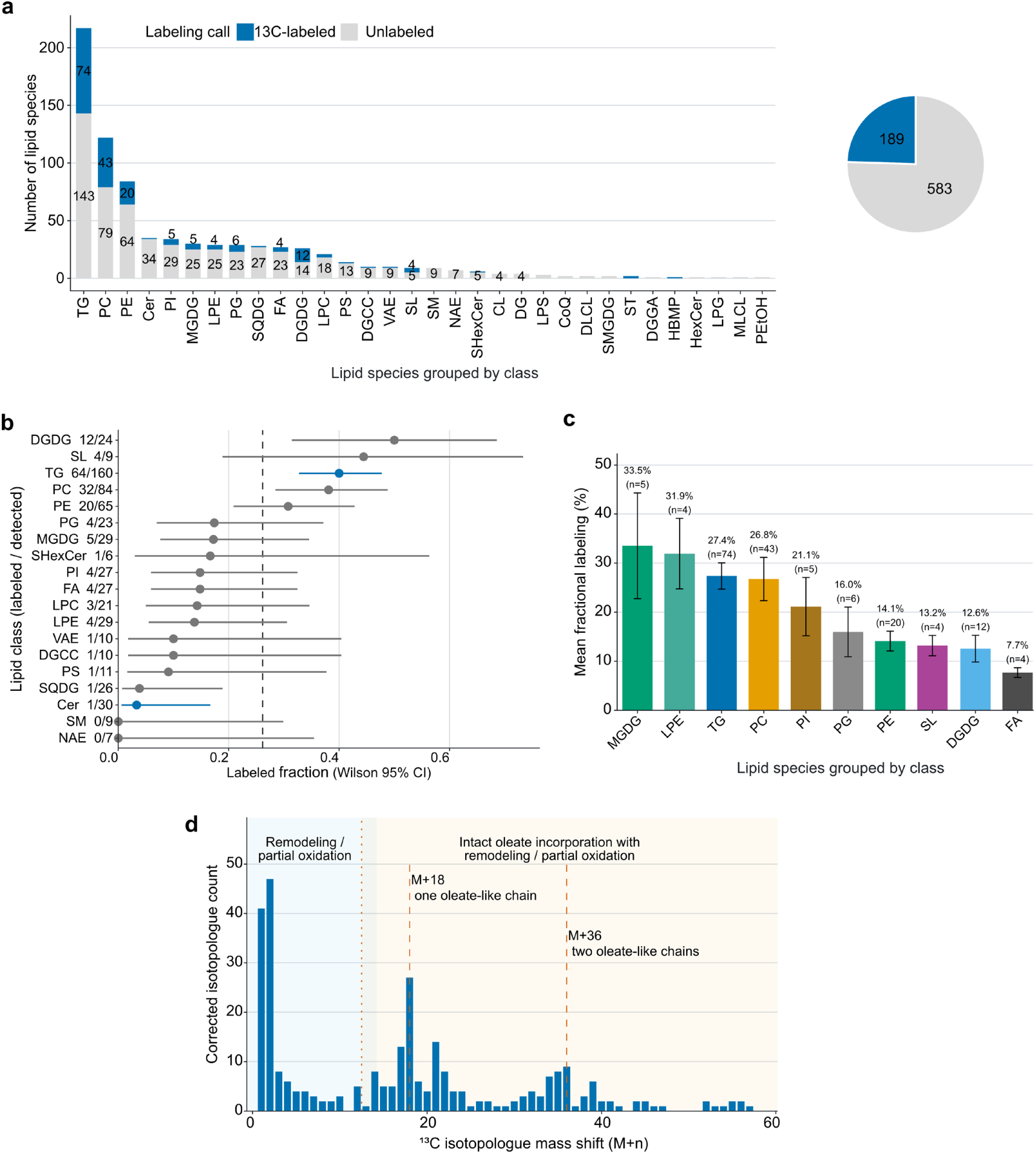
Host-acquired oleate-derived carbon enters multiple algal lipid classes. **(a)** Labeled and unlabeled structural lipid annotations by class; the inset summarizes all 772 entries. Labeling required Δ labeling > 0.05 and isotope-labeling BH-adjusted *P* < 0.05. **(b)** Class-enrichment analysis after collapse to 631 unique base/sum-composition annotations. Points show labeled fractions, lines show Wilson 95% confidence intervals, the dashed line shows the overall labeled fraction and blue denotes a class-versus-rest Fisher test with BH-adjusted *P* < 0.05. **(c)** Mean fractional labeling among labeled annotations for classes with at least four labeled entries; bars show means and error bars show SEM across annotations. **(d)** Corrected isotopologue mass-shift distribution. Dashed reference lines mark M+18 and M+36, which are consistent with incorporation of one or two oleate-derived acyl chains. Supporting Data 2, B-C; Supporting Tables S3a-S3c.

We next compared mean fractional labeling across major lipid classes to evaluate the extent of host-derived fatty acid incorporation within labeled molecules. Monogalactosyldiacylglycerol (MGDG) and LPE showed the highest mean fractional labeling, followed by TG, PC, and PI (Fig. 3c; Supporting Table S3c). These class-level patterns show that host-derived fatty-acid carbon entered both storage and membrane-lipid pools. MGDG and DGDG are major thylakoid galactolipids^34^, placing labeled host-derived carbon in photosynthetic-membrane lipids.

Examining isotopologue distributions across labeled algal lipid species allows us to assess what kind of host-derived carbon compounds are incorporated into algal lipids (see Materials and Methods for details). Class-resolved isotopologue distributions showed that the low-mass and M+18/M+36 populations occurred across multiple lipid classes, supporting broad but heterogeneous metabolic routing of oleate-derived carbon (Fig. 3d and Supporting Fig. S2d). The two higher-mass isotopologue populations (with the highest counts at M+18 and M+36) are consistent with incorporation of one or two largely intact oleate-derived acyl chains into complex algal lipids. These patterns support transfer and metabolic use of oleate-derived carbon in algal endosymbionts.

To determine whether the expanded host lipid-droplet pool could instead be explained by photosynthetically fixed carbon from the algal symbiont, we performed reciprocal isotope tracing using NaH^13^CO_3_. Symbiotic *P. bursaria* cultures were labeled with NaH^13^CO_3_ for 24 h and subsequently separated into host cytosolic, host lipid droplet-enriched, and symbiotic algal fractions for hydrophilic metabolomic and hydrophobic lipidomic analyses (Supporting Fig. S2e). Using delta labeling > 0.05 and isotope-labeling BH-adjusted *P* < 0.05, labeled features were detected in the algal metabolome (220/1,121), algal lipidome (44/624), and host cytosolic metabolome (52/1,121) (Supporting Fig. S2e; Supporting Data 2, D; Supporting Table S3d). In contrast, no host lipid-droplet lipid feature met these criteria (0/646). These results indicate that, under the conditions tested, newly fixed algal carbon contributed readily to algal metabolism and to host soluble metabolite pools, but did not yield qualifying host lipid-droplet lipid features within the present sampling window and analytical threshold. Thus, the TG-rich host lipid droplet pool observed in symbiotic *P. bursaria* is unlikely to be explained primarily by recent photosynthetic carbon fixation from the symbiont, supporting the interpretation that host lipid droplet expansion largely reflects host-side lipid remodeling.

### Lipid remodeling in *Chlorella variabilis* under symbiosis

We next examined how the symbiotic state was associated with the algal lipidome using transmission electron microscopy (TEM). After treatment with 0.1 μM oleic acid for 24 h, lipid droplets were visible in both free-living and endosymbiotic *C. variabilis*, with larger lipid-droplet profiles in the illustrated endosymbiotic cell (Fig. 4a). Cell-level quantification showed that lipid droplets occupied a greater proportion of cell area in endosymbiotic algae than in free-living algae under both untreated and oleate-treated conditions (two-sided Mann-Whitney U tests, untreated, *P* = 6.24 × 10^-9^; oleate-treated, *P* = 2.37 × 10^-10^; Fig. 4b; Supporting Data 3, A; Supporting Table S4a). This expansion of the lipid-droplet compartment suggested that symbiosis alters algal lipid accumulation.

**Figure 4.**
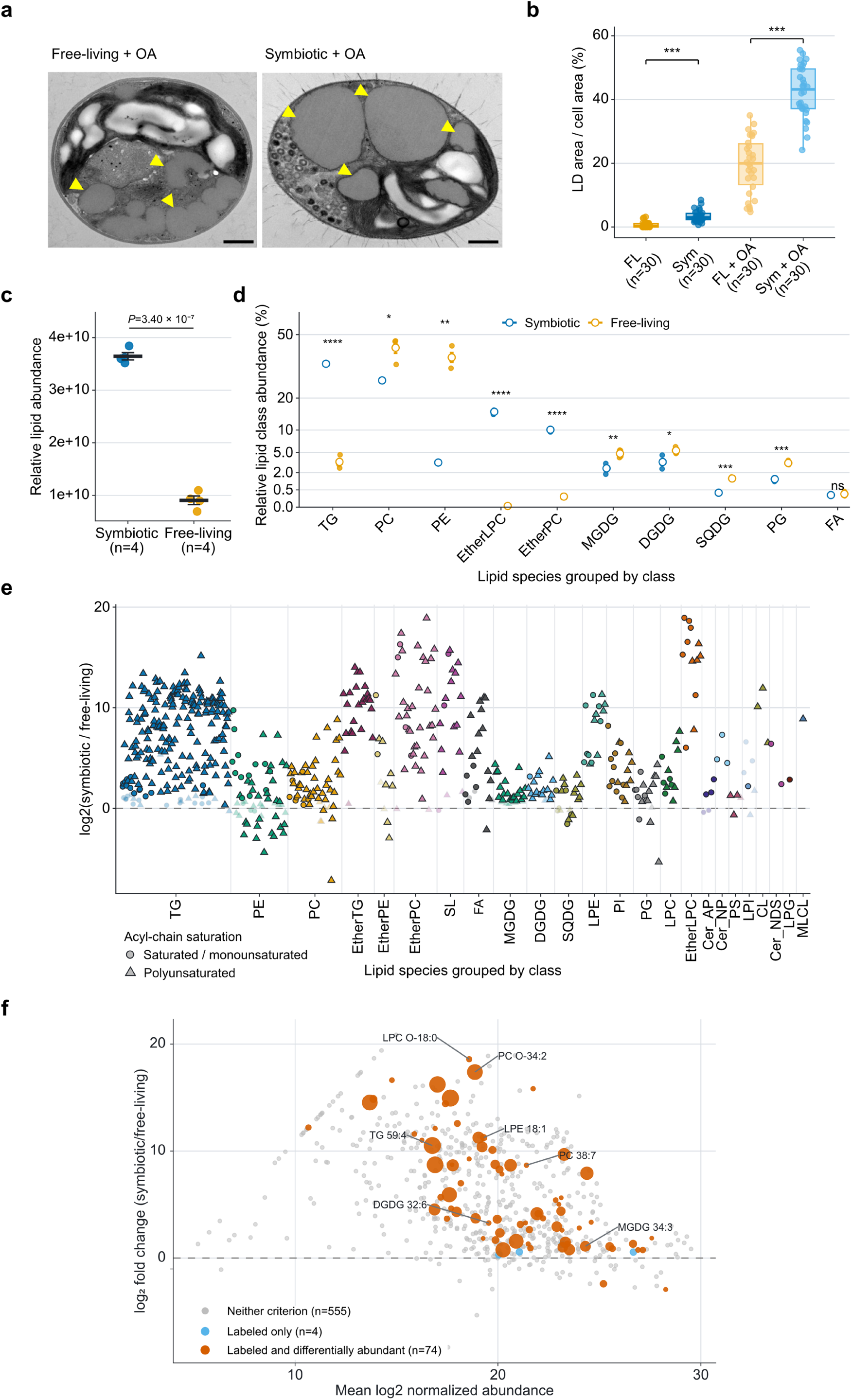
Lipid remodeling in symbiotic *Chlorella variabilis* and overlap with host-acquired oleate labeling. **(a)** TEM images of oleate-treated free-living and endosymbiotic algae. Yellow arrowheads mark lipid droplets; scale bars, 0.5 µm. **(b)** Lipid-droplet area as a percentage of cell area in untreated and oleate-treated algae (30 cells/group). Boxes show median and interquartile range; whiskers extend to 1.5 times the interquartile range. Endosymbiotic versus free-living comparisons gave *P* = 6.24 × 10^-9^ untreated and *P* = 2.37 × 10^-10^ with oleate (two-sided Mann-Whitney U tests). **(c)** Total algal lipid abundance (four biological replicates/state; mean ± SEM; 4.03-fold difference; two-sided Welch test, *P* = 3.40 × 10^-7^). **(d)** Relative abundance of ten major lipid classes (replicate points; open circles, means; error bars, SEM; square-root y axis). Asterisks mark BH-adjusted two-sided Welch tests (\**P* < 0.05, \*\**P* < 0.01, \*\*\**P* < 0.001 and \*\*\*\**P* < 0.0001; ns, *P* ≥ 0.05). **(e)** Species-level symbiotic/free-living log_2_ fold changes for 633 annotations. Fill indicates class, shape acyl-chain saturation, and opacity/outline nominal Welch *P* < 0.05 versus *P* ≥ 0.05; marker size is constant. The dashed line marks zero; x positions are categorical and compressed within each class. **(f)** MA plot of these annotations (x, mean log_2_ normalized abundance; y, symbiotic/free-living log_2_ fold change). Gray indicates neither host-fed ^13^C_18_-oleate labeling nor differential abundance, light blue labeling only, and orange-red both labeling and differential abundance (BH-adjusted *P* < 0.05; 74 annotations). Orange-red circle area scales with the tracer-minus-control difference in fractional labeling; other circles have fixed sizes. Leader lines identify seven named lipids. Supporting Data 3, A, C-D and H; Supporting Tables S4a-S4e.

To further dissect these changes at the lipidome level, we performed comprehensive LC-MS/MS profiling of endosymbiotic and free-living *C. variabilis*. Consistent with the TEM data, the total lipid content of symbiotic *C. variabilis* was approximately 4.03-fold higher than that of free-living cells (two-sided Welch test, *P* = 3.40 × 10^-7^; Fig. 4c; Supporting Table S4b). In total, 633 unique biological lipid annotations were identified and classified into 24 lipid subclasses (Supporting Fig. S3a; Supporting Table S4c). Within-condition Pearson correlations were high in free-living and symbiotic algae (r = 0.95-0.99), whereas cross-condition correlations were lower (r = 0.49-0.55), supporting replicate consistency and condition-associated lipidome remodeling (Supporting Fig. S3b; Supporting Data 3, B).

Analysis of lipid class composition showed that PC and PE were the two most dominant lipid classes in free-living cells. However, TG (34.6% of the total lipid pool) became dominant in symbiotic *C. variabilis*, a similar trend also observed in the symbiotic host cell (Fig. 4d; Supporting Data 3, C; Supporting Table S4d). Moreover, TG abundance increased by approximately 39.8-fold (two-sided Welch test on log_2_-transformed class-summed abundance, *P* = 1.91 × 10^-4^, BH-adjusted *P* = 3.82 × 10^-4^; Supporting Fig. S3c; Supporting Table S4e). Many TG species with positive fold changes contained polyunsaturated acyl chains (Fig. 4e; Supporting Data 3, D). Together, the increased lipid-droplet area, higher total lipid abundance and pronounced TG enrichment support greater neutral-lipid accumulation in symbiotic *C. variabilis*.

The class-level comparison confirmed that remodeling was unevenly distributed among neutral and membrane-associated lipid classes rather than reflecting uniform lipid accumulation (Supporting Fig. S3c). In addition to TG, several ether-linked lipids, including etherTG, etherPC, and etherLPC, were markedly increased, supporting a broader shift toward ether-lipid accumulation. At the same time, multiple membrane-associated lipid subclasses were differentially remodeled rather than uniformly elevated. Together, these patterns show compositionally selective lipid remodeling in symbiotic *C. variabilis* rather than a simple bulk increase in total lipid content.

To summarize coordinated differences in lipid composition, we used the Functional Lipid Analysis module of LipidOne^35^. LipidOne calculates indices from lipid-class abundances, acyl-chain properties and ratios between lipid pools. These indices summarize compositional features related to neutral-lipid accumulation, membrane organization and signaling-associated lipid classes. Symbiotic *C. variabilis* showed higher TG-based and neutral-to-polar lipid indices than free-living algae, consistent with the pronounced TG enrichment detected by LC-MS/MS (Supporting Fig. S4a-c; Supporting Data 3, E). Indices associated with membrane saturation, unsaturation and fluidity also differed between conditions, while the LPC/PC, LPE/PE and (LPC+LPE)/PL ratios were elevated in symbiotic algae. Acyl-chain analysis further showed lower relative representation of saturated and two-double-bond chains and greater representation of one-, three- and four-double-bond chains. Class-resolved comparisons showed that TG shifted toward longer and more unsaturated acyl chains in symbiotic algae (Supporting Fig. S4d-g; Supporting Data 3, F-G). These results extend the class-level findings by showing coordinated differences in lipid-pool distribution and acyl-chain composition.

We next compared this comparative lipidome data with the labeled lipids detected in endosymbiotic *C. variabilis* after *P. bursaria* hosts were fed ^13^C_18_-oleic acid. Their annotation identities were matched to the comparative lipidome to determine whether labeled lipids also differed in abundance between endosymbiotic and free-living *C. variabilis* (Fig. 4f; see Lipidome differential-abundance and isotope-overlap analysis in Methods). Of the 633 unique lipid annotations in the comparative lipidome, 78 matched lipids that met the isotope-labeling criteria in the host-feeding experiment (Δ labeling > 0.05 and isotope-labeling BH-adjusted *P* < 0.05). Seventy-four of these 78 annotations also differed significantly in normalized LC-MS/MS abundance between endosymbiotic and free-living algae (BH-adjusted *P* < 0.05). Among the 74 annotations, 72 had positive symbiotic-to-free-living log_2_ fold changes and two had negative log_2_ fold changes, indicating higher and lower mean abundance in endosymbiotic algae, respectively. These independent measurements therefore identified 74 algal lipid annotations that incorporated carbon from host-fed oleic acid and differed in abundance between the two algal states (Fig. 4f; Supporting Data 3, H). Their replicate-level abundance profiles showed consistent differences between conditions, supporting the reproducibility of the differential-abundance pattern (Supporting Fig. S5; Supporting Data 3, I).

Together, the imaging and lipidomics results show that endosymbiotic *C. variabilis* develops a TG-rich lipid-droplet phenotype accompanied by broader changes in lipid-class and acyl-chain composition. Many lipids that incorporated carbon from host-fed oleic acid were also more abundant in endosymbiotic than in free-living algae. Host-acquired fatty-acid carbon therefore reaches multiple algal lipid classes and overlaps extensively with the lipidomic changes associated with endosymbiosis.

### Transcriptomic remodeling of lipid metabolism in *Chlorella variabilis*

The pronounced differences in lipid abundance and composition between symbiotic and free-living *C. variabilis* prompted us to examine the corresponding transcriptional changes. We compared transcript abundance between the two algal states using RNA sequencing. Of the 8,479 transcripts retained for differential-expression analysis, 6,013 differed significantly between symbiotic and free-living algae (absolute log_2_ fold change ≥ 1 and BH-adjusted *P* < 0.05; Supporting Data 4, A-B). The transcriptome-wide distribution revealed extensive expression differences between the two algal states. Genes involved in lipid metabolism were highlighted to place their expression changes within the broader transcriptomic profile (Supporting Fig. S6; Supporting Data 4, B-C; Supporting Table S5a).

To further investigate how lipid-related pathways were influenced, we next organized the mapped genes by biochemical reaction to determine how specific lipid-metabolic processes differed between symbiotic and free-living algae. The reconstruction included fatty-acid synthesis, activation, elongation and desaturation, β-oxidation, glycerolipid synthesis, phospholipid synthesis and plastid-lipid synthesis (Fig. 5a; Supporting Data 4, C-E; Supporting Table S5b). Among the 90 unique genes displayed in this reconstruction, 46 passed the expression filter and met the differential-expression criteria. Of these, the majority of them (34/46) had higher transcript abundance in symbiotic algae. We present the measured lipid-class changes associated with the terminal reactions alongside the transcript data (Fig. 5a; Supporting Table S5c).

**Figure 5.**
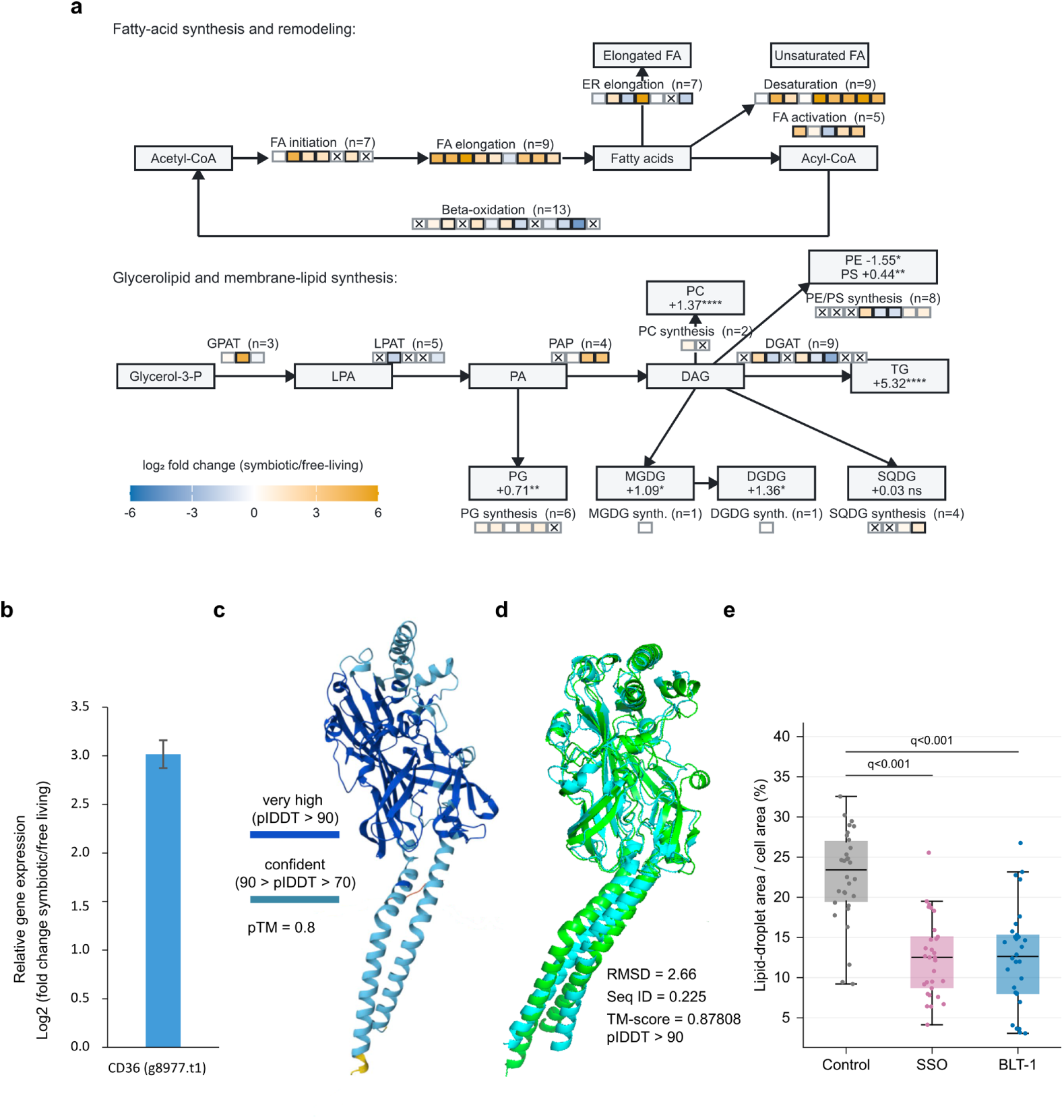
Lipid-pathway remodeling and evaluation of a CD36-like candidate in *Chlorella variabilis*. **(a)** Directed reconstruction of fatty-acid, glycerolipid, glycerophospholipid and plastid-lipid reactions. Arrows indicate reaction direction. The 93 tiles represent reaction-gene assignments from 90 unique genes; three genes occur in two reactions. Fill shows log₂ fold change for symbiotic relative to free-living algae. Black borders identify transcripts meeting both differential-expression criteria (absolute log₂ fold change ≥ 1 and BH-adjusted *P* < 0.05); gray borders indicate tested transcripts below these criteria; crosses indicate mapped transcripts below the expression filter. Terminal boxes show class-summed lipid log₂ fold changes. MGDG and DGDG synthesis are shown as sequential reactions. Asterisks indicate BH-adjusted *P* values from Welch tests across 24 lipid classes (*P* < 0.05, \**P* < 0.01, \*\**P* < 0.001 and \*\*\**P* < 0.0001; ns, *P* ≥ 0.05). **(b)** RNA-seq log₂ fold change for CD36-like candidate g8977.t1; the error bar shows propagated SE from log₂ TPM values across three biological replicates per state. **(c)** AlphaFold-predicted g8977.t1 structure colored by pLDDT confidence: dark blue, >90; cyan, 70-90; yellow, <70. The predicted template-modeling score is 0.80. **(d)** Structural superposition of g8977.t1 (green) and human CD36 (cyan); RMSD, sequence identity and TM-score are shown. **(e)** Lipid-droplet area after oleic acid alone or with SSO or BLT-1. Points represent cells (*n* = 30/group); boxes show medians and interquartile ranges, with whiskers extending to 1.5 times the interquartile range. Two-sided equal-variance Student’s *t*-tests gave BH-adjusted *P* = 2.55 × 10**^-^**⁸ for BLT-1 and 2.57 × 10**^-^**⁹ for SSO versus oleic acid alone. Supporting Data 4, C-G; Supporting Data 5, A; Supporting Tables S5b-S5c and S6a-S6d.

Together, the transcriptome-wide and pathway-resolved analyses associate the symbiotic algal state with selective remodeling of fatty-acid and complex-lipid metabolism. Increased expression of several fatty-acid synthesis and acyl-cha n-remodeling genes coincided with the accumulation of longer and more unsaturated TG species in symbiotic algae. These results identify candidate metabolic reactions that may contribute to the distinct lipid composition of endosymbiotic *C. variabilis*.

### BLT-1 and SSO reduce oleate-associated lipid-droplet accumulation

We showed that oleate-derived carbon reached endosymbiotic algae and was incorporated into the algal lipidome (Figs. 2–4). To investigate the possible mechanisms underlying algal fatty-acid uptake, we examined the genome and transcriptomes of *C. variabilis* cells. Our comparative analyses identified a CD36-like candidate (g8977.t1) with higher RNA-seq abundance in endosymbiotic algae (Fig. 5b; Supporting Table S6a), and phylogenetic analysis placed it within a broader eukaryotic CD36-like protein family that includes animal, fungal and algal sequences (Supporting Fig. S7a). CD36-family proteins are membrane-associated scavenger receptor-like proteins best characterized in animals, where CD36 participates in the recognition, binding and uptake of long-chain fatty acids and other lipid-related ligands, and has also been implicated in lipid sensing and intracellular lipid accumulation^36,37^.

Since CD36 family proteins are not well characterized in algae, we performed computational analyses to understand molecular features of this algal protein. The AlphaFold-predicted algal CD36-like protein adopts a well-defined fold with high-confidence regions (Fig. 5c). Structural comparison with human CD36 suggests substantial three-dimensional similarity despite relatively limited primary-sequence identity (Fig.^38^. Most importantly, pocket prediction identified a surface-accessible hydrophobic cavity in the algal protein (Supporting Fig. S7b) and docking analyses placed a long-chain fatty acid within a region comparable to the lipid-binding region of human CD36-related structures (Supporting Fig. S7c).

In addition to the predicted structural similarity and hydrophobic pocket, DeepTMHMM predicted that the algal CD36-family protein is membrane associated, and DeepLoc2.1 predicted plasma-membrane localization (Supporting Tables S6b and S6c^39,40^.

Guided by these computational predictions, we tested whether the oleic-acid-associated lipid-droplet phenotype was sensitive to pharmacological inhibition of scavenger-receptor lipid-uptake pathways. Block lipid transporter-1 (BLT-1) was originally characterized as an inhibitor of scavenger receptor class B type I (SR-BI)-mediated lipid transfer, whereas sulfosuccinimidyl oleate (SSO) binds mammalian CD36 and inhibits CD36-associated fatty-acid uptake^41,42^. *C. variabilis* cells were cultured in the medium with or without 2 μM BLT-1 and 2 μM SSO for 24 h, followed by 0.5 μM oleic acid feeding for 24 h. After treatments, we examined algal lipid droplet size using TEM. BLT-1- and SSO-treated cells exhibited significantly lower lipid-droplet area than untreated cells (two-sided equal-variance Student t-tests; BLT-1, BH-adjusted *P* = 2.55 × 10^-8^; SSO, BH-adjusted *P* = 2.57 × 10^-9^; Fig. 5e). This inhibitor-sensitive phenotype strongly supports the possible role of the scavenger-receptor pathway in algal lipid uptake (Supporting Data 5, A; Supporting Table S6d). Nonetheless, the uptake efficiency and substrate specificity remain to be determined.

## Discussion

Photosymbioses are commonly described as partnerships in which photosynthetic symbionts export fixed carbon to heterotrophic hosts, while hosts provide inorganic nutrients and a protected intracellular habitat^4,5^. Classic studies of *Paramecium bursaria* established algal-to-host photosynthate transfer and host provisioning of nitrogenous resources^43,44^. Our findings add a complementary direction. Under oleate supplementation, carbon acquired by the host reached intracellular *Chlorella variabilis* and entered algal storage and membrane lipids. Together with established photosynthate export, this result supports reciprocal movement of organic carbon at the partnership level. Reciprocity here refers to bidirectional exchange through chemically distinct carbon pools whose magnitudes and timing may differ.

Lipid exchange is increasingly recognized as a component of symbiotic integration, with mechanisms shaped by each partnership^11,45^. The enlarged triacylglycerol-rich host lipid-droplet pool provides a cellular setting for this exchange. Lipid droplets are dynamic metabolic organelles that respond to diverse physiological states and exert important regulatory functions^46,47^. This view agrees with our earlier finding that the triacylglycerol biosynthetic pathway is upregulated in the host cell and disrupting host triacylglycerol metabolism reduces symbiont abundance^25^. Moreover, reciprocal NaH^13^CO_3_ tracing yielded little evidence for rapid contribution of newly fixed algal carbon to the expanded host neutral-lipid pool. NanoSIMS and EA-IRMS showed that carbon from host-acquired ^13^C_18_-oleic acid reached intracellular algae. Isotopologue-resolved lipidomics then localized this carbon within algal lipids. M+18 and M+36 species supported incorporation of one or two largely intact oleate-derived acyl chains, whereas low-mass isotopologues were consistent with degradation and recycling. This mixture indicates two distinct fates for host-derived carbon: partial incorporation as intact acyl chains through algal re-esterification, and partial degradation and recycling.

Transferred carbon entered both neutral-storage and membrane-lipid pools. Labeling of TG establishes access to algal neutral-lipid storage, whereas phospholipid, MGDG, and DGDG labeling places host-derived carbon in membrane-associated lipid synthesis. MGDG and DGDG form a major part of the thylakoid lipid matrix^48,49^. The high incorporation frequency among DGDG species therefore links transferred acyl carbon to a lipid class central to photosynthetic membranes. TGs can serve as reversible buffers that connect acyl sequestration, membrane remodeling, and stress recovery in green algae and plants^50,51^. These patterns support a hypothesis in which host-derived acyl carbon expands a flexible substrate reserve that can be divided between storage and membrane maintenance. Consistently, several genes assigned to fatty-acid synthesis, elongation, and desaturation had higher transcript abundance in symbiotic algae.

A central biological question is why a photosynthetic symbiont would receive organic carbon from its host. One possibility is metabolic division of labor (DOL), in which partners distribute metabolic functions and resource demands through metabolite exchange^52,53^. Feeding-derived fatty acids provide concentrated carbon and preassembled acyl chains. Symbiont access to this pool could reduce some biosynthetic demand for producing fatty acids de novo during transient light limitation or intracellular resource constraint. This possibility is consistent with evidence that host phagotrophy benefits intracellular *Chlorella* by providing access to particulate nutrient sources^54^. Stable algal metabolism could in turn promote symbiont persistence and preserve the future return of photosynthate to the host. Host provisioning may therefore contribute to partner stability, with benefits emerging over a longer interval than the initial carbon-transfer event.

Photoautotrophy can coexist with the use of environmental organic carbon. In green hydra, glucose uptake by some *Chlorella* symbionts was associated with persistence in darkness, and host feeding stimulated algal division^55^. The *C. variabilis* NC64A genome also retains broad metabolic capacity^56^. In the same host system studied here, symbiotic *Micractinium conductrix* displayed higher carbon fixation together with a marked increase in lipid-droplet volume^57^. These observations show that photosynthetic productivity, organic-carbon use and lipid storage can coexist in hospite. Host-derived fatty acids may therefore complement photosynthesis rather than substitute for it.

## Materials and Methods

### Cell culture conditions

*P. bursaria* cells were cultured in CGM medium (12 mM sodium acetate, 0.5% yeast extract. 2 mM sodium citrate, 1 mM NaH_2_PO_4_/Na_2_HPO_4_, and 1.5 mM CaCl_2_). Cultures were maintained at 23 °C under a 12 h light/12 h dark photoperiod and were fed with *Chlorogonium capillatum* cells every five days.

Aposymbiotic *P. bursaria* strains were generated by treating symbiotic cells with 100 μg/ml of methyl viologen dichloride hydrate (Paraquat dichloride, Cat #856177, Sigma-Aldrich, MA, USA) for approximately one week in a light-to-dark cycle of 12 h:12 h^58^. The resulting aposymbiotic cells were subsequently maintained under low-light conditions.

Free-living *Chlorella variabilis* cells were cultivated in modified CA medium. The medium was prepared by mixing 1 mL of 2% Ca(NO_3_)_2_·4H_2_O, 1 mL of 10% KNO_3_, 1 mL of 5% NH_4_NO_3_, 1 mL of 3% β-Na_2_-glycerophosphate·5H_2_O, 1 mL of 2% MgSO_4_·7H_2_O, 10 mL of 4% HEPES buffer, 1 mL of PIV metals solution, and 10 mL of Fe solution (as EDTA, 1:1 molar ratio). Vitamin supplements were added as follows: 10 μL of 1 mg mL-¹ vitamin B₁₂ (final concentration 10 μg mL-¹), 100 μL of 0.1 mg mL-¹ biotin (final concentration 1 μg mL-¹), and 10 μL of 1 mg mL-¹ thiamine hydrochloride. The final volume was adjusted to 1 L using double-distilled water (ddH₂O), and the pH was adjusted to 7.2. The PIV metals solution was prepared by combining 10 mL of 1% Na_2_EDTA·2H_2_O, 1.96 mL of 1% FeCl_3_·6H_2_O, 0.36 mL of 1% MnCl_2_·4H_2_O, 0.11 mL of 2% ZnSO_4_·7H_2_O, 0.10 mL of 0.4% CoCl_2_·6H_2_O, and 0.10 mL of 0.25% Na_2_MoO_4_·2H_2_O, followed by adjustment of the total volume to 100 mL with ddH_2_O. The Fe-EDTA solution was prepared by mixing 7.02 mL of 1% Fe(NH_4_)_2_(SO_4_)_2_·6H_2_O with 6.6 mL of 1% Na_2_EDTA·2H_2_O and adjusting the final volume to 100 mL with ddH_2_O.

### Lipid droplets isolation and metabolites extraction

We collected four biological replicates for each *P. bursaria* sample. For each replicate of lipid extraction from lipid droplet samples, approximately 1 × 10^5^ symbiotic or aposymbiotic *P. bursaria* cells (corresponding to ∼1 L of culture volume) were harvested through a 10 μm nylon mesh sieve after removing cell debris with cheesecloth filtration. The collected cells were washed on the nylon mesh with Dryl’s solution multiple times and subsequently transferred into 15 mL centrifuge tubes. The washed cells were resuspended in lysis buffer (10 mM MgCl_2_, 10 mM Tris pH 7.5, and 250 mM sucrose) and lysed using a nitrogen cavitation bomb. The bomb was pressurized to 300 psi and maintained for 20 minutes, after which the pressure was slowly released to lyse the cells. Following cell disruption, the lysate was mixed with an equal volume of high sucrose buffer (10 mM MgCl_2_, 10 mM Tris pH 7.5, and 2.1 M sucrose) and centrifuged at 17,000 rpm for 1 hour at 4 °C using an SW41Ti rotor (Beckman Coulter, CA, USA). The topmost lipid layer was carefully collected and further purified by re-floating the lipid droplets through centrifugation at 13,000 × g for 10 minutes at 4 °C. The upper lipid droplet fraction was then collected for downstream lipid extraction analysis. For hydrophilic metabolite extraction from *P. bursaria*, the cytosolic fraction obtained following cell disruption.

For lipids and hydrophilic metabolite extraction from *Chlorella* cells, approximately 2 × 10^6^ free-living *Chlorella variabilis* cells were harvested by centrifugation at 8,000 × g, followed by two washes with Dryl’s buffer. A similar number of symbiotic *Chlorella* cells were isolated from *P. bursaria* by cell disruption through high-speed centrifugation at 20,000 × g for 20 minutes, followed by debris removal using a 10 μm nylon mesh and two washes with Dryl’s buffer. We collected four biological replicates for each *Chlorella* sample.

Lipid extraction was performed following a modified chloroform/methanol method^59^. Isolated lipid droplets were resuspended to final volume about 500 µL in lysis buffer, while *Chlorella* cells were resuspended in 500 µL of Dryl’s buffer. Subsequently, 1 mL of pre-cooled chloroform/methanol (2:1, v/v) was added to each sample. The mixtures were gently agitated for 1 hour on ice to ensure complete lipid extraction. After incubation, 0.25 mL of cold double-distilled water (ddH₂O) was added, and the samples were vortexed for 10 minutes while kept on ice. Phase separation was achieved by centrifugation at 13,000 rpm for 10 minutes at 4 °C. The upper hydrophilic layer was carefully discarded, and the lower hydrophobic (organic) phase was transferred to a new microcentrifuge tube, taking care to avoid disturbing the interfacial protein disc. The collected organic phase was dried using a SpeedVac concentrator and stored at -80 °C until further analysis. Prior to LC-MS/MS analysis, the dried lipid extracts were reconstituted in 80 µL of chloroform/methanol (2:1, v/v) containing a lipid isotope internal standard mixture (Avanti Research, Cat # A83707, AL, USA). The samples were then centrifuged at 13,000 rpm for 10 minutes at 4 °C, and the resulting supernatant was collected for LC-MS/MS analysis. Hydrophilic metabolites were extracted using the same chloroform/methanol-based extraction procedure described below for lipid extraction. Following phase separation, however, the upper hydrophilic phase was carefully collected instead of being discarded. The collected hydrophilic phase was dried using a SpeedVac concentrator and stored at -80 °C until further analysis. Prior to LC-MS/MS analysis, the dried hydrophilic metabolite extracts were reconstituted in water/methanol (1:1, v/v), centrifuged at 13,000 rpm for 10 minutes at 4 °C, and the resulting supernatants were collected for metabolomic analysis.

### Lipidomic LC-MS/MS analysis

For each biological replicate sample, we performed four technical replicate experiments. Lipidomic profiling was conducted utilizing an Orbitrap Fusion Lumos Tribrid MS (Thermo Fisher Scientific, MA, USA) coupled with a Vanquish Horizon UHPLC system (Thermo Fisher) equipped with a BEH C18 column (100 × 2.1 mm, 1.7 µm; Waters). Lipid separation was achieved through a 14 min gradient employing 40% acetonitrile and 10% ACN/90% isopropanol with 10 mM ammonium formate at a flowrate of 0.5 mL/min. Both MSs were operated in a data-dependent acquisition mode (DDA) featuring one MS scan and multiple MS/MS scans at a fixed cycle time of 0.4 sec. MS/MS spectra were acquired under collision-induced dissociation (CID, 25 and 35 eV) and higher-energy collisional dissociation (HCD, normalized collision energy 25 and 45, stepped ±15%). Raw MS data were processed using MS-DIAL software (version 4.90)^31^. The parameters for peak detection were set as follows: minimum peak height 30000 amplitude, minimum peak width of 10 scans, and peak smoothing using a linear weighted moving average over four scans. Metabolite identification was performed using a combined MS/MS spectral library composed of the NIST Tandem Mass Spectral Library and the MassBank of North America (MoNA: https://mona.fiehnlab.ucdavis.edu/). Lipid identification was performed using the LipidBlast MS/MS library, supplemented with in-house predicted retention times. Mass tolerances were set to 0.008 Da for MS1 and 0.05 Da for MS/MS, with a retention-time tolerance of 0.5 min for lipids. A score cutoff of 75 was applied.

Hydrophilic extracts were analyzed using an Orbitrap IQ-X Tribrid mass spectrometer (Thermo Fisher Scientific, Waltham, MA) coupled to a Vanquish UHPLC system (Thermo Fisher Scientific). Data were acquired in positive and negative electrospray-ionization modes. Chromatographic separation used either a HILIC-Z column (2.1 × 100 mm, 2.7 µm; Agilent, Santa Clara, CA) or an HSS T3 column (2.1 × 100 mm, 1.8 µm; Waters, Milford, MA). For HILIC-Z separation, the mobile phases contained acetonitrile and ammonium formate (pH 3) in positive mode or ammonium acetate (pH 9) in negative mode; a 12-min gradient was applied at 0.5 mL/min. For HSS T3 separation, the mobile phases were 2% acetonitrile with 0.1% formic acid and acetonitrile with 0.1% formic acid; a 10-min gradient was applied at 0.4 mL/min. Lipidomic profiling used an Orbitrap Fusion Lumos Tribrid mass spectrometer coupled to a Vanquish Horizon UHPLC system and a BEH C18 column (100 × 2.1 mm, 1.7 µm; Waters). Lipids were separated over a 14-min gradient using 40% acetonitrile and 10% acetonitrile/90% isopropanol containing 10 mM ammonium formate at 0.5 mL/min. Both mass spectrometers were operated in data-dependent acquisition mode with one MS1 survey scan followed by multiple MS/MS scans in a fixed 0.4-s cycle. MS/MS spectra were acquired using collision-induced dissociation (25 and 35 eV) and higher-energy collisional dissociation (normalized collision energies of 25 and 45 with stepped ±15%).

Raw MS data were processed using MS-DIAL version 4.90^31^. Peak-detection parameters were a minimum peak height of 30,000, a minimum peak width of 10 scans and linear-weighted moving-average smoothing over four scans. Metabolites were identified against a combined MS/MS spectral library comprising the NIST Tandem Mass Spectral Library and MassBank of North America. Lipids were identified using the LipidBlast MS/MS library supplemented with in-house predicted retention times^60^. Mass tolerances were 0.008 Da for MS1 and 0.05 Da for MS/MS, the lipid retention-time tolerance was 0.5 min and the score cutoff was 75. Isotope-tracing data were processed using El-MAVEN^32^.

### TEM and SEM imaging

Symbiotic *P. bursaria* cells were incubated for 24 h with 0.1 mM unlabeled or uniformly ^13^C-labeled oleic acid for isotope-tracing experiments. Endosymbiotic and free-living *Chlorella* cells were incubated for 24 h with 0.1 μM oleic acid, with parallel untreated controls used for lipid-droplet comparisons. Samples were washed repeatedly before NanoSIMS, EA-IRMS or electron-microscopy preparation. For TEM and SEM, samples were fixed in 2.5% glutaraldehyde in 0.05 M cacodylate buffer (pH 7.2) using a PELCO 3451 laboratory microwave system (Ted Pella, CA) for one cycle of 1 min on, 1 min off and 1 min on at 250 W; a ColdSpot module maintained the samples at 10 °C. After rinsing in 0.05 M cacodylate buffer, samples were postfixed with 1% OsO4 in the same buffer using one cycle of 1 min on, 1 min off, 1 min on and 1 min off at 100 W. Samples were rinsed with double-distilled water, dehydrated through a graded ethanol series using one cycle of 1 min on, 1 min off and 1 min on at 100 W, infiltrated with a graded Spurr resin series for 3 min at each step at 250 W, embedded in plastic capsules and polymerized at 70 °C for 12 h. Ultrathin sections of approximately 100 nm were cut using a Leica EM UC6/UC7 ultramicrotome. TEM images were acquired using a Tecnai G2 Spirit TWIN transmission electron microscope (Thermo Fisher Scientific) operated at 120 kV and equipped with a Gatan SC1000 CCD camera (4,008 × 2,672 active pixels). A single-tilt holder, CompuStage rotation holder or Gatan 626 cryo specimen holder was used as appropriate.

SEM images of thin sections were acquired using an FEI Helios NanoLab 660 (Thermo Fisher Scientific) in high-resolution mode with a circular backscatter detector. The beam current was 0.2 nA, the dwell time was 30 μs and the working distance was 4 mm. A stage bias of -1.5 kV was applied to improve the signal-to-noise ratio. The primary beam energy was 4.5 keV, giving a landing energy of 3.0 keV. Signals from all four detector segments were summed, and image contrast was inverted to provide TEM-like contrast.

### NanoSIMS imaging

Carbon-isotope ratios in thin sections of symbiotic *P. bursaria* cells incubated with oleic acid were mapped using a NanoSIMS 50L instrument (Cameca-Ametek, Gennevilliers, France) at Academia Sinica, Taiwan. Microtomed sections were mounted on 10-mm-diameter silicon wafers. Each 50 × 50 μm region of interest was presputtered with a 165-172 pA Cs+ ion beam for 1 min and then sputtered with a 2.8-2.9 pA Cs+ ion beam for 45 min. Images were acquired at 512 × 512 pixels. Secondary ions of ^16^O-, ^12^C_2_-, ^12^C^13^C-, ^12^C^14^N-, ^31^P- and ^32^S-were collected simultaneously using electron multipliers. A 30-μm entrance slit and a 150-μm aperture slit provided a mass-resolving power (M/ΔM) of 10,000. Secondary-ion and carbon-isotope-ratio images were processed using L’Image software (Larry Nittler, Arizona State University). More than 10⁵ counts of ^12^C_2_-were collected from each algal region.

### EA-IRMS analysis and Stable isotope tracing

For EA-IRMS analysis, endosymbiotic algae were isolated from symbiotic *P. bursaria* cells fed with oleic acid using high-speed centrifugation (21,000 × g, 10 min). The collected algal cells were washed with 1X phosphate buffered saline (PBS) buffer multiple times to remove *P. bursaria* debris completely. Algal cells were dried and weighed to the required mass, then enclosed in tin capsules for subsequent analysis. During the analysis, international isotope ratio standards USGS40, USGS62, and USGS65 (United States Geological Survey Reston Stable Isotope Laboratory: https://www.usgs.gov/labs/reston-stable-isotope-laboratory) were inserted as reference materials for calibration and calculation. The isotope ratio analysis was performed using an EA-Conflo-IRMS system. Samples were combusted into gas form by the Thermo Flash2000 elemental analyzer (Thermo Fisher Scientific), transferred through a Conflo IV interface (Thermo Fisher Scientific), and analyzed using the Thermo Delta V Advantage isotope ratio mass spectrometer (Thermo Fisher Scientific). The EA system was connected to two high-purity gases: oxygen (5N), which assisted the combustion of the samples, and helium (5N), which served as the carrier gas. When the sample dropped into the combustion tube heated to 1050 °C, it was oxidized to produce gaseous compounds including NO_x_, CO_2_, SO_2_, and H_2_O. SO_2_ was absorbed by silver-cobalt oxide packed within the combustion tube. The remaining gases (NO_x_, CO_2_, and H_2_O) then entered the reduction tube, where they were converted into N_2_, CO_2_, and H_2_O. Subsequently, the gas stream was passed through a water-removal column containing magnesium perchlorate, which absorbed H₂O, leaving only N_2_ and CO_2_. These purified gases were then introduced into the isotope ratio mass spectrometer, where the isotopic ratios of N_2_ and CO_2_ were precisely measured.

For isotope-tracing LC-MS/MS, symbiotic *P. bursaria* cells were incubated for 24 h with 0.1 mM unlabeled or uniformly ^13^C-labeled oleic acid (^13^C_18_-oleic acid; MedChemExpress, HY-N1446S2) or with 100 mg/L NaH^12^CO_3_ or NaH^13^CO_3_ (MedChemExpress, HY-W251598S). Metabolites and lipids were extracted and analyzed by LC-MS/MS as described above. Isotopologue peaks were extracted with El-MAVEN, and natural-isotope abundance was corrected using IsoCorrectoR^33^. Assignment of the ^13^C label to specific acyl-chain positions was validated by MS/MS.

Isotopologue peak intensities corresponding to M, M+1, up to M+N were extracted for each metabolite from the LC-MS/MS datasets. Quantification of ^13^C incorporation was performed by summing the intensities of all labeled isotopologues (M+1 through M+N), yielding the total labeled signal. To normalize for variations in metabolite abundance across samples, the fractional ^13^C labeling was calculated as the ratio of the summed labeled isotopologues to the total isotopologue signal (M to M+N).

For downstream comparisons, fractional labeling values were stratified into two experimental groups: ^13^C-labeled and control. For each metabolite, the mean fractional labeling of the ^13^C-labeled group was contrasted with that of the control group. The difference between these means (Δ labeling = mean_^13^C - mean_control) was used as the metric of ^13^C enrichment. Statistical evaluation of group-wise differences was conducted independently for each metabolite by Student’s t-test. A metabolite was designated as significantly ^13^C-labeled if it satisfied both of the following criteria: (1) the mean fractional labeling in the ^13^C-labeled group exceeded that of the control group (Δ labeling > 0.05); and (2) the Benjamini-Hochberg-adjusted *P* value was < 0.05.

### *C. variabilis* genome assembly and annotation

Because the available *C. variabilis* reference assembly remained fragmented^56^, a strain-specific reference genome was assembled and annotated using published reads together with newly generated sequencing data. For genome assembly, Illumina HiSeq 4000 paired-end sequencing reads and MinION long-read sequencing data for *C. variabilis* NC64A (ATCC 50528) were obtained from NCBI BioProject PRJDB7392^61^. Read quality was evaluated using FastQC (v0.11.5) for short reads and NanoQC (v0.9.4) for long reads^62^. Low-quality reads were trimmed using Trimmomatic (v0.39) ^63^for short reads and NanoFilt (v2.8.0) ^62^for long reads. Hybrid genome assembly was performed using MaSuRCA (v4.1.0) ^64^with default parameters. Assembly quality was evaluated using QUAST (v5.2.0)^65^. The assembly was further refined using Redundans (v2.0.1)^66^ under scaffolding conditions with long-read support.

Variants between NIES-2540 and NC64A were identified using HaplotypeCaller implemented in GATK^67^, and the resulting variants were incorporated into the NC64A reference genome to generate the NIES-2540 strain-specific genome. The final genome assembly consisted of 260 contigs with a total size of 44.2 Mb and a GC content of 67.1%. The N50 value of the assembly was 296,995 bp. RNA-seq reads were mapped to the assembled genome using HISAT2 (v2.2.1)^68^, yielding an alignment rate of approximately 88%. Gene structure prediction was performed using BRAKER3 (v3.0.8)^69^, resulting in 14,574 predicted transcripts. Genome completeness was assessed using BUSCO (v5.7.1)^70^ with the Eukaryota odb10 dataset (255 single-copy orthologs), showing approximately 93% completeness.

### RNA isolation and sequencing

For RNA isolation, approximately 3 × 10^8^ algal cells were collected from freshly isolated symbiotic algae and long-term free-living cultures. Cells were washed twice with 1× PBS and resuspended in nuclease-free water to a final volume of 200 μL. All samples were prepared in triplicate. For RNA extraction, 0.8 mL TRI reagent and 0.5 mL of 0.01 mm glass beads were added to each sample. Cells were homogenized using a bead beater (FastPrep-24 5G Instrument; MP Biomedicals, Irvine, CA) for 80 s at room temperature, then immediately cooled on ice. Lysates were centrifuged at 12,000 × g for 10 min at 4 °C to remove debris. The supernatant was transferred, mixed with 50 μL of 0.5 M potassium acetate, and incubated for 5 min at room temperature. Subsequently, 300 μL chloroform was added and mixed vigorously for 3 min. Phase separation was achieved by centrifugation at 21,000 × g for 20 min at 4 °C. The aqueous phase was transferred to 0.6 mL of ice-cold 100% ethanol. RNA purification was subsequently performed using the RNeasy Mini Kit (Qiagen). RNA quality was assessed prior to library preparation. Strand-specific RNA libraries were prepared using the SureSelect Strand-Specific RNA Library Kit and sequenced on a NovaSeq X Plus platform. Approximately 20 million paired-end reads were generated per sample.

### Transcriptome analysis

Raw sequencing reads were evaluated using FastQC (v0.11.5) (http://www.bioinformatics.babraham.ac.uk/projects/fastqc/), and adapter sequences and low-quality reads were removed using Trimmomatic^63^. Transcript abundance was quantified with Salmon (v2.7.0^71^. Transcripts were retained when TPM was at least 1 in at least two of three biological replicates in either condition. Differential expression was tested with limma using log_2_(TPM + 0.1) values^72^. *P* values were adjusted using the Benjamini-Hochberg procedure. Differential expression was defined by an absolute log_2_ fold change ≥ 1 and BH-adjusted *P* < 0.05.

### Functional annotation and lipid-pathway reconstruction

Predicted *C. variabilis* proteins were functionally annotated using complementary sequence-based approaches. Protein domains and Gene Ontology assignments were obtained using InterProScan ^73^ and Blast2GO^74^. KEGG Orthology identifiers were assigned using GhostKOALA and DeepKOALA, and assignments from the two analyses were combined^75,76^. MapMan4 functional categories were assigned using Mercator4 version 8.0^77^. The resulting gene, GO, KEGG Orthology and MapMan annotations were assembled into an organism-level annotation database using AnnotationForge^78^. The resulting gene descriptions, Gene Ontology terms, KEGG Orthology mappings, and MapMan assignments were consolidated for downstream analysis.

Genes associated with lipid metabolism were identified from their KEGG Orthology and MapMan4 assignments. KEGG module definitions were used to organize orthologous enzymes into fatty-acid synthesis, activation, elongation and degradation, glycerolipid, glycerophospholipid and plastid-lipid reactions^79,80^. Mercator4 assignments supplemented reactions that were not adequately represented by the selected KEGG modules, including fatty-acid desaturation and galactolipid synthesis. All genes assigned to each reaction were retained, including multiple paralogs. Salmon-derived TPM values and limma differential-expression results were then mapped to the corresponding reactions.

### Lipid compositional index analysis using LipidOne

Annotated lipid species from symbiotic and free-living *Chlorella variabilis* were analyzed using LipidOne (https://lipidone.eu/)^35^. Lipid-property indices were calculated based on lipid class, acyl-chain length, and degree of unsaturation, and were compared between conditions using log_2_ fold change and statistical significance. Lipids were also grouped by unsaturation number, and summed peak areas within each group were compared between symbiotic and free-living algae. LipidOne supplied raw *P* values for 29 Energy, Structural and Signaling indices; Benjamini-Hochberg adjustment was applied across all displayed indices. Composition-normalized unsaturation profiles and class-resolved abundance-weighted chain-length and double-bond summaries were calculated from the biological-replicate lipid matrix.

### Confocal microscopy

For lipid droplet staining, cells were similarly fixed with 4% formaldehyde for 10 minutes and stained with LipidSpot^TM^ dye (1000X dilution; Cat. # 70065-T, Biotium, CA, USA) for 30 minutes at room temperature. Fluorescence signals were detected using a ZEISS LSM 780 upright confocal microscope (ZEISS, Oberkochen, Germany), with an excitation wavelength of 440 nm and an emission wavelength of 585 nm.

### Phylogenetic analysis of CD36-like proteins

The predicted *Chlorella variabilis* CD36-like protein sequence was used as a query to retrieve homologous sequences from UniProt BLAST results^81^. Amino acid sequences of CD36-like proteins were collected and aligned using MAFFT (v7) with default parameters^82^. Poorly aligned or ambiguous regions were inspected and adjusted when necessary. Phylogenetic relationships were inferred by maximum likelihood using IQ-TREE (v2)^83^. The best-fit amino acid substitution model was selected automatically by IQ-TREE, and branch support was evaluated by bootstrap analysis. The resulting tree was visualized and annotated with taxonomic information using Interactive Tree of Life (iTOL, v7.6)^84^.

### Protein structure prediction and structural comparison

The three-dimensional structure of the *C. variabilis* CD36-like candidate was predicted using AlphaFold 3^85^. To assess structural similarity between the algal candidate and known CD36-family proteins, pairwise structural comparison was performed using TM-align^86^. The resulting TM-score, RMSD, and sequence identity values were used as summary metrics to evaluate structural similarity between the predicted algal CD36-like protein and the reference CD36 structure.

### Pocket prediction and docking analysis

Potential ligand-binding pockets in the predicted algal CD36-like structure were analyzed using CASTpFold^87^. To test whether the predicted structure could accommodate long-chain fatty acids, docking analysis was performed using AutoDock Vina ^88^ with palmitic acid (FA 16:0) ^89^as the ligand. Docking poses were visualized using PyMOL ^90^ and UCSF Chimera^91^. The predicted fatty-acid-binding mode of the algal CD36-like protein was compared with the corresponding ligand-binding region in the human CD36 structure.

### DeepLOC and DeepTMHMM predictions

For subcellular localization, protein feature annotations were predicted using DeepLoc 2.1^40^. Transmembrane spans were predicted in the annotated *Chlorella variabilis* protein sequences using DeepTMHMM^39^.

### Lipidome differential-abundance and isotope-overlap analysis

To determine whether incorporation of host-fed oleate-derived carbon occurred among the *C. variabilis* lipids whose abundance differed between symbiotic and free-living states, we compared results from the comparative lipidomics and host-feeding isotope-tracing experiments. The comparative dataset comprised lipid-abundance measurements from symbiotic and free-living *C. variabilis*, with four biological replicates per condition. The isotope-tracing dataset comprised labeling measurements from endosymbiotic *C. variabilis* isolated after their hosts were fed ^13^C_18_-oleic acid. Lipid names were standardized across the two datasets, duplicate rows corresponding to the same biological lipid annotation were collapsed and annotations were matched by exact identity. For each annotation in the comparative dataset, mean log_2_ abundance and the symbiotic-to-free-living log_2_ fold change were calculated. Differences between conditions were evaluated using two-sided Welch tests followed by Benjamini-Hochberg correction. Isotope labeling was defined as Δ labeling > 0.05 and isotope-labeling BH-adjusted *P* < 0.05. Annotations that met the isotope-labeling criterion and differed significantly between symbiotic and free-living algae were classified as supported by both datasets (BH-adjusted *P* < 0.05). The matched results were displayed in an MA plot, with point size among annotations supported by both datasets representing Δ labeling. Replicate abundance profiles for these annotations were visualized using row-standardized values.

### Pharmacological inhibition assay

To test whether oleate-associated lipid-droplet accumulation was sensitive to pharmacological probes of scavenger-receptor lipid-uptake pathways, inhibitor experiments were performed using 2 μM BLT-1 (Cat. No.: HY-116767, MedChemExpress), originally characterized as an SR-BI inhibitor, and 2 μM sulfosuccinimidyl oleate sodium (SSO) (Cat. No.: HY-112847A, MedChemExpress), an inhibitor of mammalian CD36-associated fatty-acid uptake. Cells were first pretreated with the indicated inhibitor for 24 h, followed by incubation with 0.5 μM oleic acid for an additional 24 h in the continued presence of the inhibitor. After treatment, samples were prepared for transmission electron microscopy (TEM). The effect of inhibitor treatment on oleate-associated lipid-droplet accumulation was assessed from lipid-droplet area and ultrastructural changes in TEM images.

### Statistical analysis

Group-comparison tests were two-sided unless otherwise stated. Biological-replicate total lipid abundances were compared using Welch t tests on untransformed summed signals. Absolute lipid-class fold changes were tested using Welch t tests on log_2_(class-summed signal + 1), followed by Benjamini-Hochberg correction across all tested classes. Relative lipid-class percentages were compared using Welch t tests with Benjamini-Hochberg correction across the displayed classes. Species-level lipid abundances were compared using Welch t tests, with Benjamini-Hochberg correction across species. NanoSIMS measurements and algal lipid-droplet area comparisons under untreated or oleate-treated conditions were evaluated using Mann-Whitney U tests, with individual analyzed cells treated as experimental units. EA-IRMS measurements were compared using equal-variance Student t tests. Inhibitor measurements were compared with equal-variance Student t tests, and *P* values were adjusted across the two prespecified comparisons with the oleate-only control. PCA separation was evaluated by exact-label PERMANOVA. Exact *P* values are reported throughout.

### Use of generative artificial intelligence

Codex (OpenAI; GPT 5.6, accessed in 2026) was used to assist with the organization, documentation, debugging, verification, and refinement of data-analysis and plot-generation code. AI assistance was also used to draft or revise code for data processing, plotting, file-format conversion, and assembly of the reproducible analysis pipeline. The authors inspected, tested, and executed all code, and verified all outputs against the underlying experimental data and final manuscript figures. The authors retain full responsibility for the analyses and their reproducibility.

## Data availability

The mass-spectrometry lipidomic data have been deposited in Metabolomics Workbench under accession PR003131 (http://dx.doi.org/10.21228/M8ZP0G). Processed data underlying the analyses are provided in Supporting Data 1-5 and Supporting Tables S1-S6. The *C. variabilis* genome assembly, annotation, and transcriptomic sequencing data have been submitted to NCBI (BioProject PRJNA1518359); the accession will be added upon release (Reviewer-only link: https://dataview.ncbi.nlm.nih.gov/object/PRJNA1518359?reviewer=ojnrkq086tocokqv9blgavuh6m). Analysis code used to generate quantitative figures, and statistical outputs have been submitted to Figshare (Reviewer only link: https://figshare.com/s/055c3dce5efa47cdeda5).

## Acknowledgements

We thank members of the Leu lab for helpful discussion and comments on the manuscript. We also thank the Academia Sinica IMB Imaging Core for instructing confocal and TEM imaging experiments, Dr. Chia-Wei Lee at Academia Sinica Volume EM & Tomography Core Facility (ASVET) for SEM imaging experiments and Academia Sinica Metabolomics Core Facility for lipidomics experiments. This work was supported by Academia Sinica of Taiwan (grant no. AS-IA-110-L01 and AS-GCS-113-L03) and the National Science and Technology Council of Taiwan (NSTC 115-2311-B-001-031-MY3). MMK was supported by an NSTC postdoctoral fellowship (NSTC 113-2811-B-001-065).

## Author contribution

JYL conceived the study. YJC, MMK and JYL designed the analysis and interpreted the results. YJC, CYW, CWH, SYH, CLY and PLW performed experiments. YJC, CYW, MMK, CLY, CWH, SYH, DCL and PLW performed data analysis and conducted a content review of the manuscript. JYL and CK provided resources. YJC, MMK and JYL wrote the paper. All authors read and approved the final manuscript.

## Declaration

The authors declare no competing interests.

