## Supplementary Data for "Host-to-symbiont transfer of fatty-acid-derived carbon in the *Paramecium bursaria*-*Chlorella variabilis* photosymbiosis"

Chen Y.J. et al.

*Correspondence

Md Mostafa Kamal; Jun-Yi Leu

**This PDF contains**

**Supporting figures S1 to S7**

**Supporting tables S1 to S6**

**Supporting Data 1 to 5 captions**

### Supporting Figures

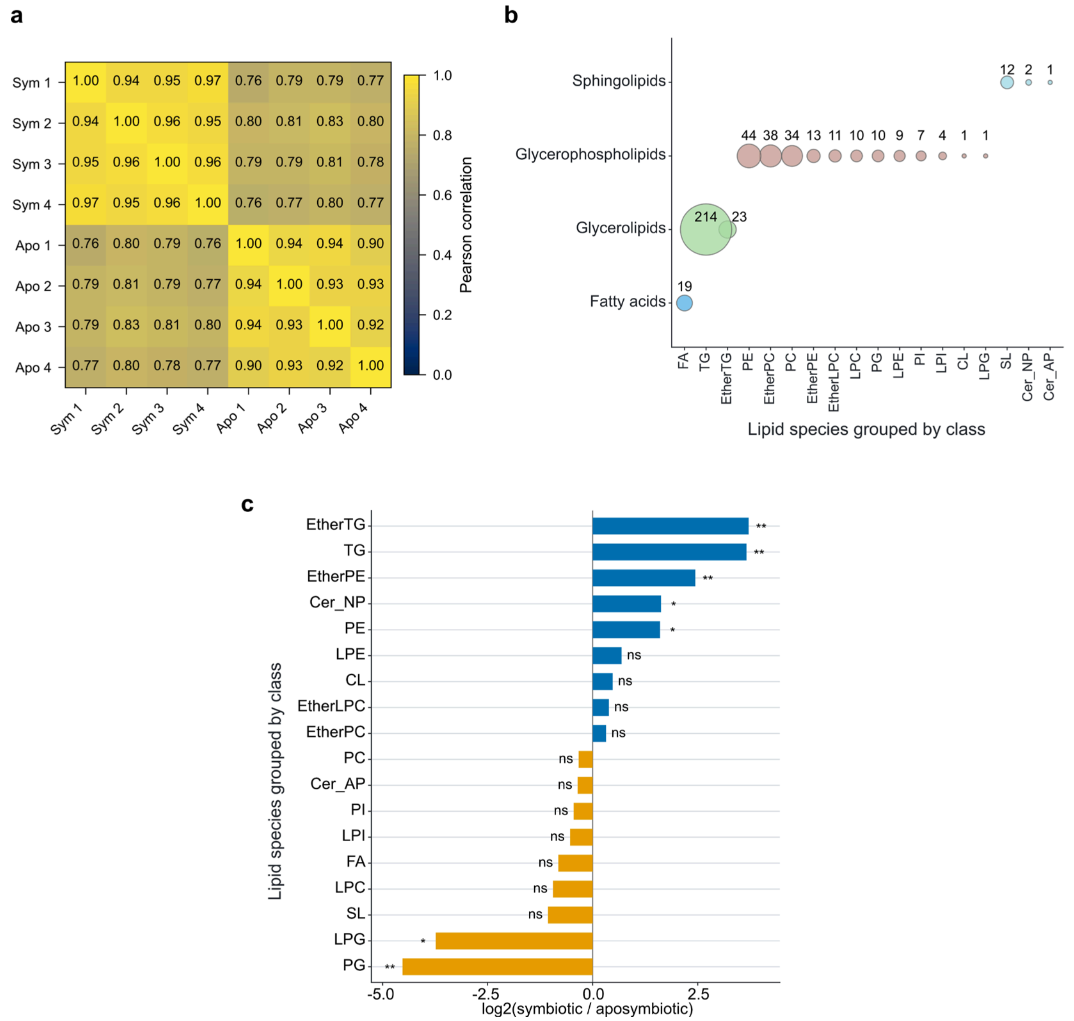


**Supporting Figure S1. Host lipidome replicate correlations, fold-change-analysis coverage and class-level differential abundance.**

**(a)** Pearson correlation matrix for host biological replicates. Cell values and fill color show Pearson r. **(b)** Numbers of host lipid annotations eligible for fold-change analysis by class. Bubble area and printed values show annotation counts; 453 annotations across 18 classes had nonzero mean abundance in both states. PS and Cer_NDS were detected only in the 478-annotation raw dataset but were excluded from fold-change analysis because one state had zero mean abundance. **(c)** Host lipid-class $\log_2$ fold changes calculated from class-summed abundance. Asterisks denote BH-adjusted two-sided Welch tests on $\log_2$(signal + 1) (*$P < 0.05$, **$P < 0.01$, ***$P < 0.001$ and ****$P < 0.0001$; ns, $P \geq 0.05$). Supporting Data 1, A and C; Supporting Tables S1c-S1d.

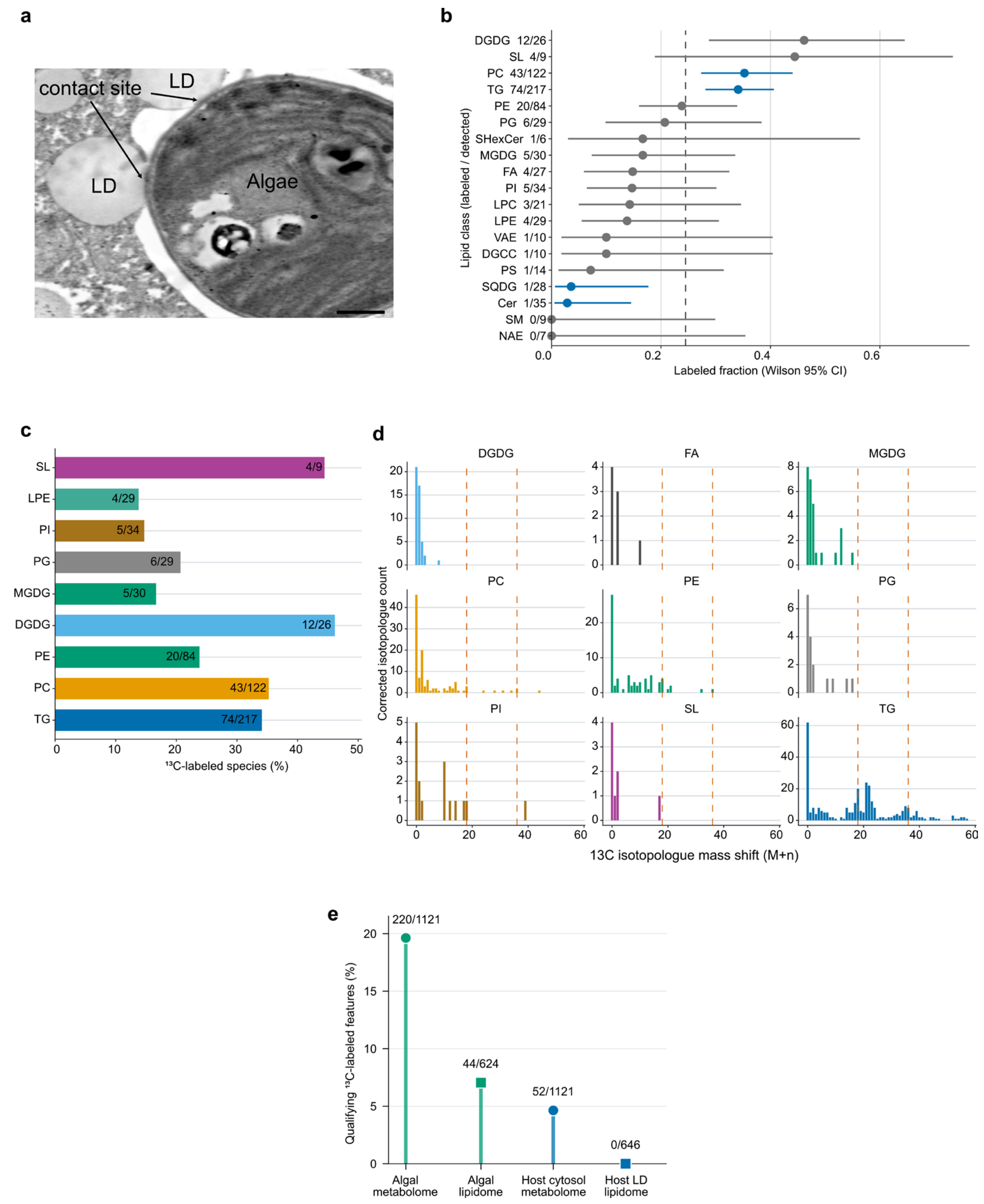


**Supporting Figure S2. Spatial context and complementary isotope-tracing analyses of host-symbiont carbon exchange.**

**(a)** Backscatter SEM image of a thin section showing a host lipid droplet adjacent to an intracellular alga; contrast is inverted to resemble a TEM image, and the scale bar represents 0.5 µm. **(b)** Structural-entry class enrichment among the 772 lipid annotations analyzed after host feeding with $^{13}C_{18}$-oleic acid. Points show labeled fractions, lines show Wilson 95% confidence intervals, the dashed line shows the overall labeled fraction and blue identifies classes with classversus-rest Fisher BH-adjusted $P < 0.05$. **(c)** Fraction labeled for the nine classes with at least four labeled entries; labels show labeled/detected annotations. **(d)** Corrected isotopologue mass-shift distributions for the same nine classes, including SL; orange dashed lines mark M+18 and M+36. **(e)** Features satisfying the $NaH^{13}CO_3$-tracing criteria after 24 h relative to $NaH^{12}CO_3$ controls. Numerators are qualifying labeled features and denominators are statistically testable features; features lacking a calculable adjusted *P* value are excluded from the denominators. Supporting Data 2, B-D; Supporting Tables S3a, S3b and S3d.

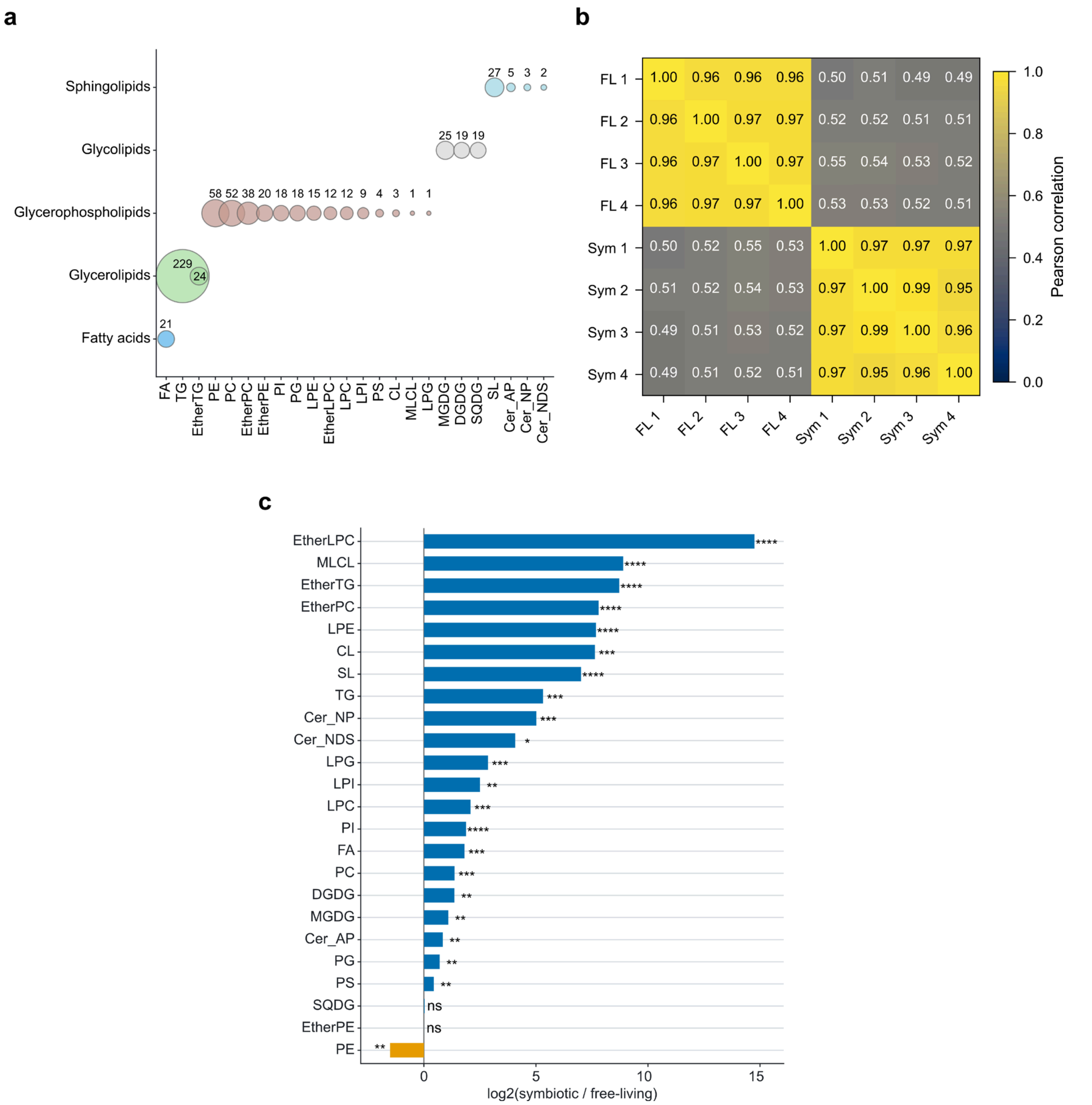


**Supporting Figure S3. Algal lipidome class coverage, replicate correlations and differential abundance.**

**(a)** Detected algal lipid annotations by class. Bubble area and printed values show annotation counts; 633 unique annotations were detected across 24 classes. **(b)** Pearson correlation matrix for algal biological replicates. Cell values and fill color show Pearson r. **(c)** Algal lipid-class $\log_2$ fold changes calculated from class-summed abundance. Asterisks denote BH-adjusted two-sided Welch tests on $\log_2$(signal + 1) (*$P < 0.05$, **$P < 0.01$, ***$P < 0.001$ and ****$P < 0.0001$; ns, $P \geq 0.05$). Supporting Data 3, B and D; Supporting Tables S4c-S4e.

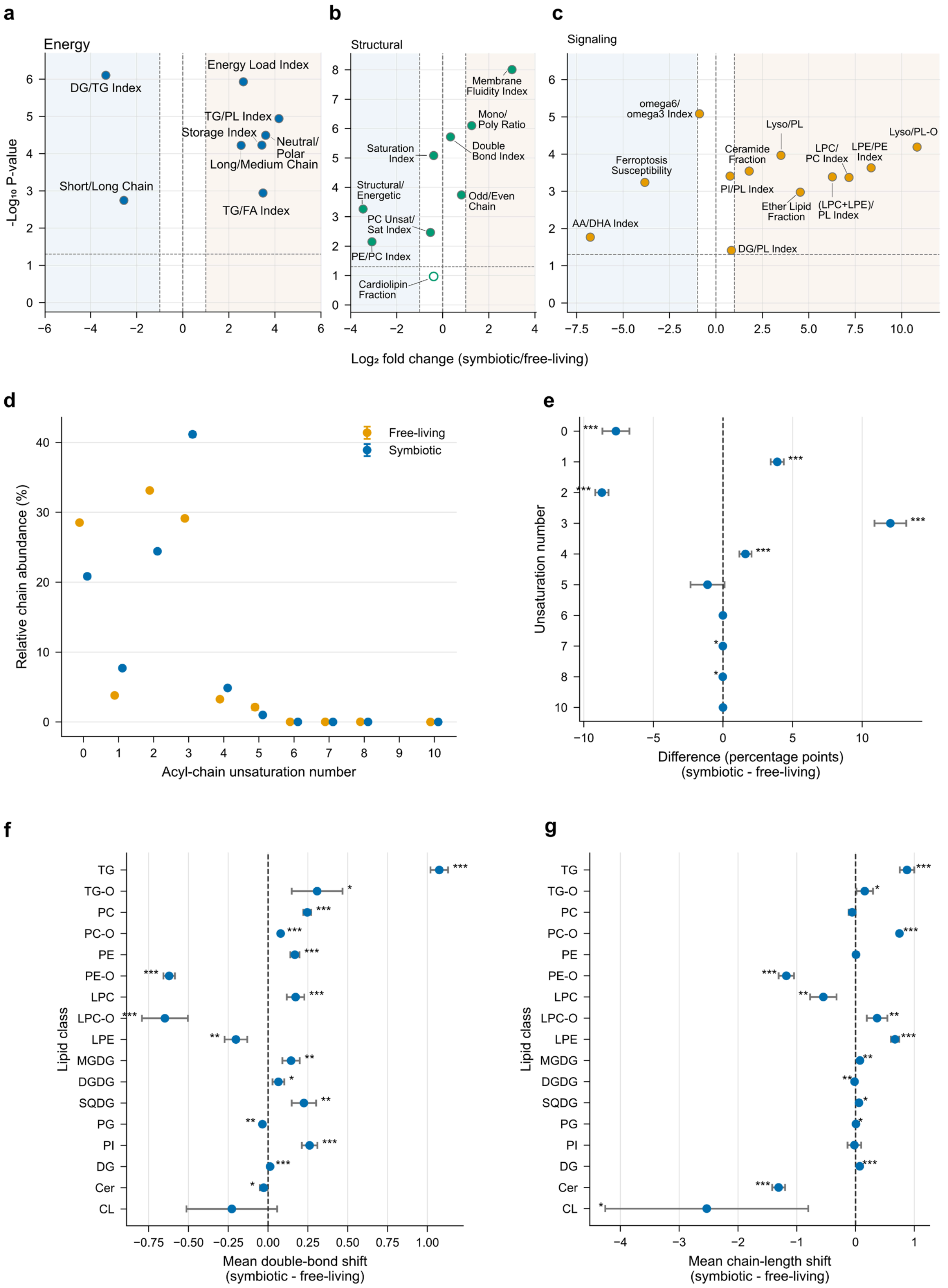


**Supporting Figure S4. LipidOne-derived compositional indices and acyl-chain remodeling**

**in symbiotic versus free-living algae.**

**(a-c)** Complete LipidOne Energy, Structural and Signaling index outputs. The x axis shows the symbiotic/free-living $log_2$ fold change and the y axis shows -log10 raw *P*. Filled points have BH-adjusted $P < 0.05$ across all 29 indices; open points do not. **(d)** Composition-normalized acyl-chain-unsaturation profiles. Points show condition means and error bars show SEM across four biological replicates. **(e)** Symbiotic-minus-free-living differences in unsaturation-bin percentages with 95% confidence intervals. **(f)** Class-resolved differences in abundance-weighted mean double-bond number with 95% confidence intervals. **(g)** Class-resolved differences in abundance-weighted mean chain length with 95% confidence intervals. Panels d-g use two-sided Welch tests with BH correction within each analysis family. Supporting Data 3, E-G.

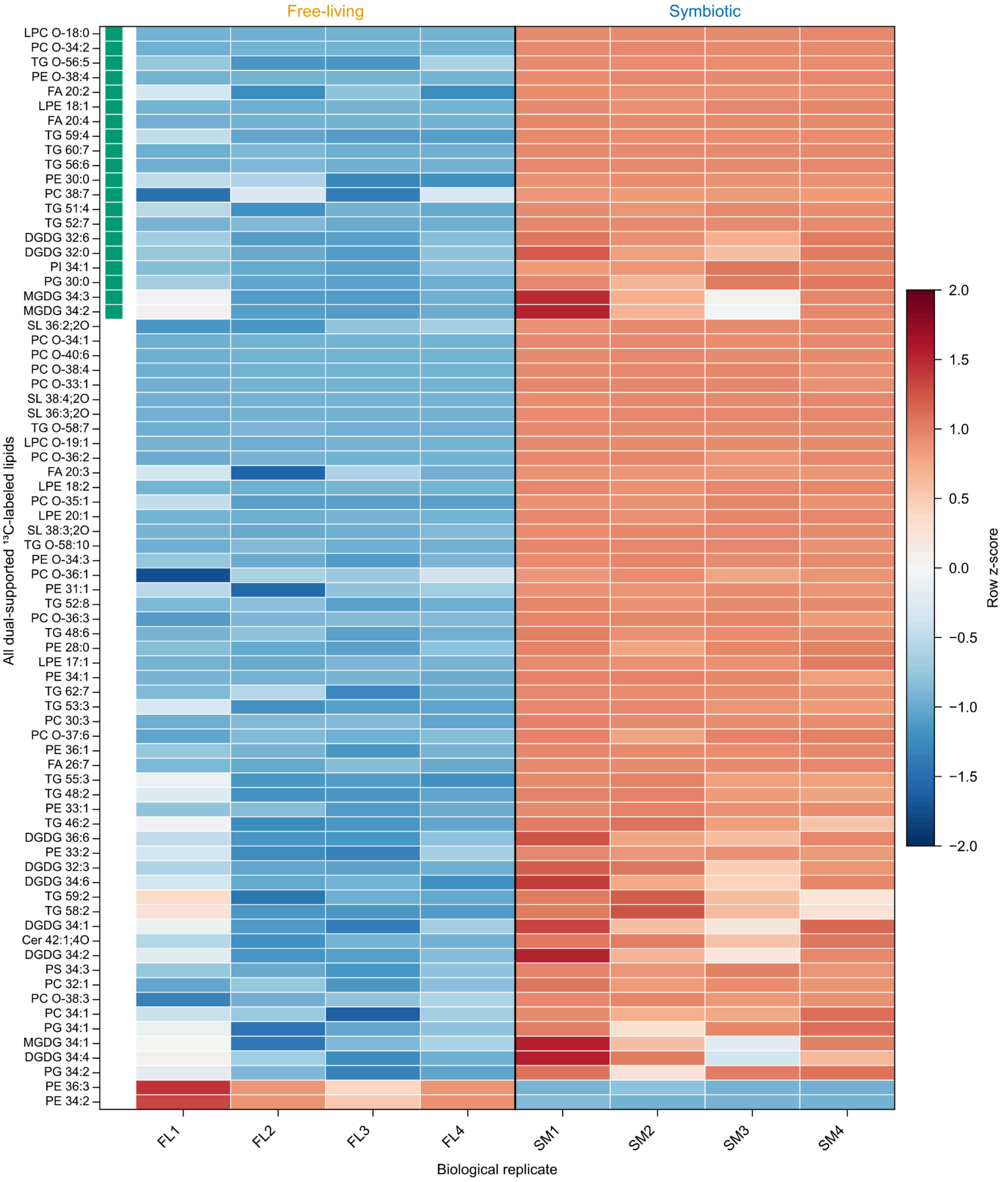


**Supporting Figure S5. Replicate abundance profiles of algal lipids supported by isotope labeling and differential abundance.**

Heatmap of all 74 lipid annotations that were both $^{13}$C labeled after host oleate feeding and differentially abundant in the symbiotic algal lipidome. The first 20 rows are green-marked manuscript-relevant representatives; the remaining 54 follow the same effect-size ordering. Row-wise z scores show within-lipid relative abundance and do not compare absolute abundance among lipids. Supporting Data 3, I.

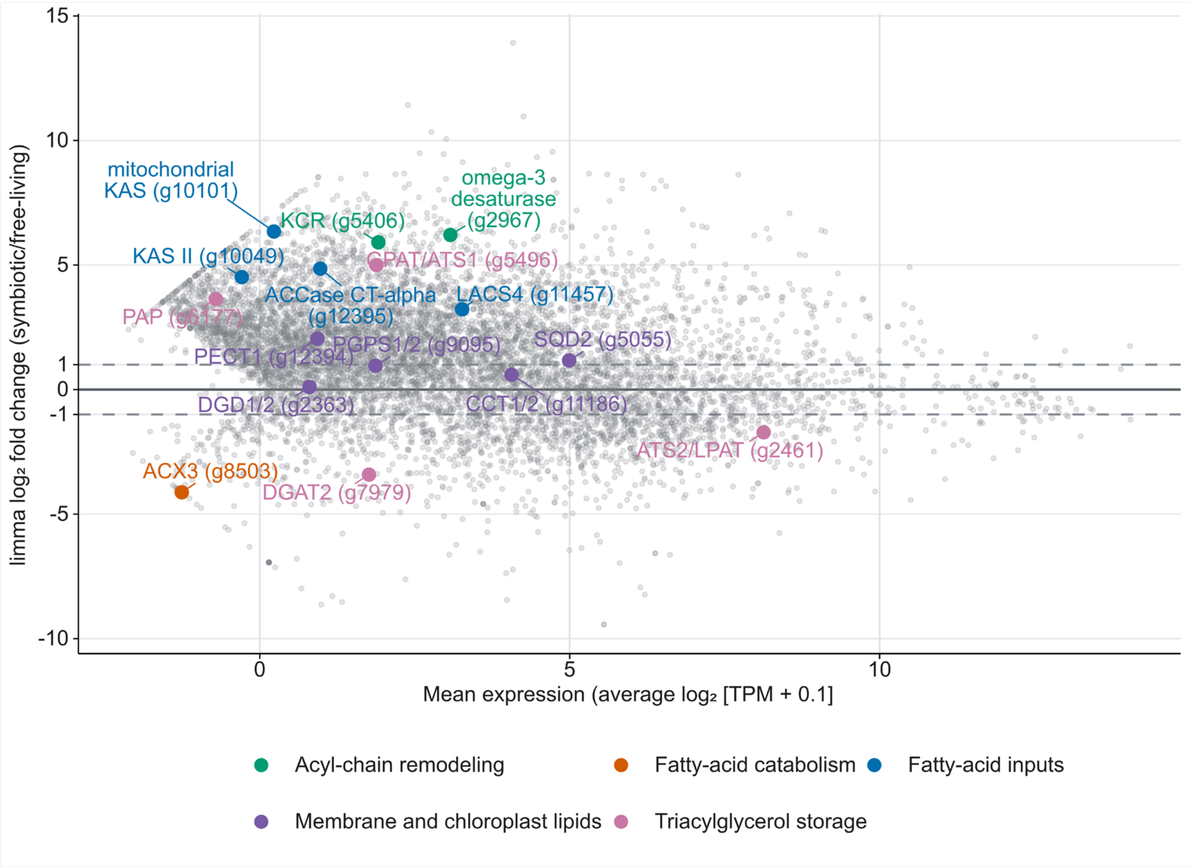


**Supporting Figure S6. Transcriptome-wide differential expression and selected lipid-pathway genes in *Chlorella variabilis*.**

MA plot of the 8,479 transcripts retained for differential-expression analysis. The x axis shows mean expression as average $log_2$(TPM + 0.1), and the y axis shows the limma $log_2$ fold change for symbiotic relative to free-living algae. Gray circles represent all tested transcripts. Colored, labeled circles identify 16 representative lipid-pathway genes grouped by functional category. The solid horizontal line marks zero fold change, and dashed lines mark $log_2$ fold changes of +1 and -1. Supporting Data 4, B-C; Supporting Table S5a.

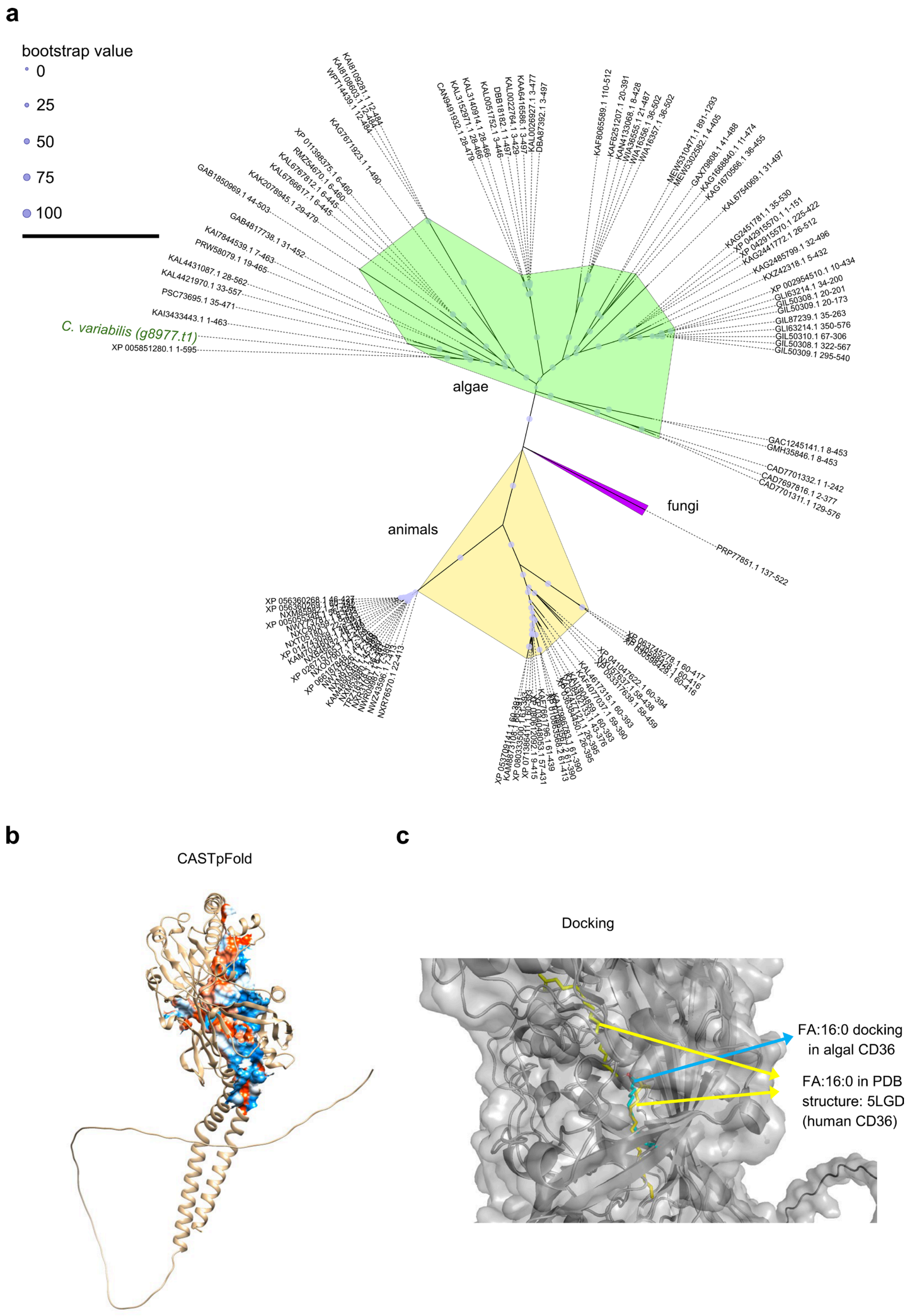

a
bootstrap value
0
25
50
75
100
C. variabilis (g8977.t1)
algae
animals
fungi
b
CASTpFold
c
Docking
FA:16:0 docking in algal CD36
FA:16:0 in PDB structure: 5LGD (human CD36)

**Supporting Figure S7. Evolutionary relationships, predicted pocket and fatty-acid docking of the *C. variabilis* CD36-like candidate.**

**(a)** Maximum-likelihood phylogeny of CD36-like proteins across major eukaryotic lineages. Node-circle size denotes bootstrap support, and the *C. variabilis* candidate g8977.t1 is highlighted in green. **(b)** CASTpFold pocket analysis of the predicted g8977.t1 structure. Surface patches are colored by Kyte-Doolittle hydrophobicity from blue (most hydrophilic) through white to red (most hydrophobic). **(c)** Palmitate docking comparison. Cyan shows the palmitate pose docked in the predicted algal CD36-like structure, and yellow shows palmitate in the human CD36 reference structure (PDB 5LGD).

**Supporting Tables S1-S6: Statistical and Analytical Summaries**

**Table S1. Host lipidome abundance, composition and class-level statistics**

Supporting-data source: Supporting Data 1, A-C

Host lipid-droplet-enriched fractions from symbiotic and aposymbiotic cells are compared using total lipid signal, lipid-class relative abundance and species-level effects. Relative-abundance tests and absolute-signal fold changes are reported separately to distinguish compositional shifts from changes in lipid-pool abundance.

**Table S1a. Total normalized host lipid-droplet signal in individual symbiotic and aposymbiotic biological replicates**

The eight signals underlying Figure 1b are shown in arbitrary units. Figure 1b compares group means using a two-sided Welch test (8.15-fold; $P = 0.0269$).

| Group | Biological replicate | Total normalized lipid signal (a.u.) |
|---|---|---|
| Symbiotic | G-LD_1 | 27344923877.8438 |
| Symbiotic | G-LD_2 | 11224761986.3852 |
| Symbiotic | G-LD_3 | 16096549437.9286 |
| Symbiotic | G-LD_4 | 30844791219.9972 |
| Aposymbiotic | W-LD_1 | 2141585731.09604 |
| Aposymbiotic | W-LD_2 | 2846895570.80228 |
| Aposymbiotic | W-LD_3 | 2843979777.51472 |
| Aposymbiotic | W-LD_4 | 2660328918.52181 |

**Table S1b. Welch tests of host lipid-class relative abundance in symbiotic versus aposymbiotic host cells**

| Lipid class | n (symbiotic) | n (aposymbiotic) | *P* value | BH-adjusted *P* | Significance |
|---|---|---|---|---|---|
| EtherLPC | 4 | 4 | 0.002307 | 0.008447 | ** |
| EtherPC | 4 | 4 | 0.007682 | 0.01097 | * |
| EtherTG | 4 | 4 | $3.180 \times 10^{-4}$ | 0.00318 | ** |
| FA | 4 | 4 | 0.003923 | 0.008447 | ** |
| LPC | 4 | 4 | 0.01002 | 0.01252 | * |
| LPE | 4 | 4 | 0.01843 | 0.02048 | * |
| PC | 4 | 4 | 0.005068 | 0.008447 | ** |
| PE | 4 | 4 | 0.05973 | 0.05973 | ns |
| SL | 4 | 4 | 0.002537 | 0.008447 | ** |
| TG | 4 | 4 | 0.004655 | 0.008447 | ** |

**Table S1c. Numbers of host lipid annotations eligible for fold-change analysis by lipid class**

| Lipid class | Analyzed species |
|---|---|
| TG | 214 |
| PE | 44 |
| EtherPC | 38 |
| PC | 34 |
| EtherTG | 23 |
| FA | 19 |
| SL | 12 |
| EtherPE | 13 |
| PG | 10 |
| EtherLPC | 11 |
| LPC | 10 |
| PI | 7 |
| LPE | 9 |
| LPI | 4 |
| Cer_NP | 2 |
| Cer_AP | 1 |

| | |
|---|---|
| CL | 1 |
| LPG | 1 |

**Table S1d. Host lipid-class absolute-signal effects expressed as $\log_2$ (symbiotic/aposymbiotic)**

| Lipid class | $\log_2$FC (symbiotic/aposymbiotic) | *P* value | BH-adjusted *P* | Significance |
|---|---|---|---|---|
| Cer_AP | -0.3569 | 0.5914 | 0.5914 | ns |
| Cer_NP | 1.628 | 0.006430 | 0.02315 | * |
| CL | 0.4753 | 0.5268 | 0.5578 | ns |
| EtherLPC | 0.3864 | 0.3344 | 0.4299 | ns |
| EtherPC | 0.3169 | 0.4780 | 0.5378 | ns |
| EtherPE | 2.443 | 0.001348 | 0.006066 | ** |
| EtherTG | 3.713 | $8.790 \times 10^{-4}$ | 0.006066 | ** |
| FA | -0.8165 | 0.08594 | 0.1547 | ns |
| LPC | -0.9444 | 0.04516 | 0.1016 | ns |
| LPE | 0.6883 | 0.1455 | 0.2381 | ns |
| LPG | -3.736 | 0.009082 | 0.02725 | * |
| LPI | -0.5381 | 0.2160 | 0.3239 | ns |
| PC | -0.3354 | 0.3597 | 0.4317 | ns |
| PE | 1.604 | 0.01083 | 0.02785 | * |
| PG | -4.525 | $5.710 \times 10^{-4}$ | 0.006066 | ** |
| PI | -0.4517 | 0.2925 | 0.4050 | ns |
| SL | -1.062 | 0.06328 | 0.1266 | ns |
| TG | 3.662 | 0.001228 | 0.006066 | ** |

Significance codes reflect BH-adjusted *P* (q-value): ns, $P \geq 0.05$; *$P < 0.05$; **$P < 0.01$; ***$P < 0.001$; ****$P < 0.0001$.

**Table S2. NanoSIMS and EA-IRMS summaries of host-derived $^{13}C$ transfer to intracellular algae**

**Table S2a. NanoSIMS and EA-IRMS enrichment summaries after host oleate feeding**

Supporting-data source: Supporting Data 2, A; individual EA-IRMS values in Table S2b.

Cell-level NanoSIMS $^{13}C/^{12}C$ percentages and biological-replicate EA-IRMS $^{13}C$ atom percentages after hosts were fed $^{13}C$-oleate, $^{12}C$-oleate or no oleate. Higher enrichment after $^{13}C$-oleate feeding quantifies host-acquired oleate carbon recovered in intracellular algae.

| Analysis | Group | n | Mean (%) | SD (%) | SEM (%) |
|---|---|---|---|---|---|
| NanoSIMS | $^{12}C$ oleic acid | 30 | 2.129 | 0.05845 | 0.01067 |
| NanoSIMS | $^{13}C$ oleic acid | 30 | 4.016 | 0.3945 | 0.07203 |
| EA-IRMS | $^{12}C$ oleic acid | 4 | 1.075 | $5.000 \times 10^{-4}$ | $2.500 \times 10^{-4}$ |
| EA-IRMS | $^{13}C$ oleic acid | 4 | 4.6 | 0.2974 | 0.1487 |
| EA-IRMS | Control | 4 | 1.08 | 0.001708 | $8.539 \times 10^{-4}$ |

**Table S2b. Biological-replicate EA-IRMS 13C atom percentages after host oleate feeding**

EA-IRMS 13C atom percentages for the three host-feeding conditions shown in Figure 2e.

| Host-feeding condition | Biological replicate | 13C atom (%) |
|---|---|---|
| Control | 1 | 1.078 |
| Control | 2 | 1.079 |
| Control | 3 | 1.08 |
| Control | 4 | 1.082 |
| 12C oleic acid | 1 | 1.075 |
| 12C oleic acid | 2 | 1.076 |
| 12C oleic acid | 3 | 1.075 |
| 12C oleic acid | 4 | 1.075 |
| 13C oleic acid | 1 | 4.68 |
| 13C oleic acid | 2 | 4.9 |
| 13C oleic acid | 3 | 4.63 |
| 13C oleic acid | 4 | 4.19 |

**Table S3. Oleate- and bicarbonate-isotope-labeling summaries and annotation-unit sensitivity**

Supporting-data source: Supporting Data 2, B-D

Isotope-labeling summaries from host-fed $^{13}$C-oleate and NaH$^{13}$CO$_3$ tracing. Features were classified as labeled when delta labeling (control minus $^{13}$C condition) exceeded 0.05 and the BH-adjusted *P* value was <0.05; features without a calculable adjusted *P* value were classified as indeterminate. The tables report lipid-class labeling, compartment-level classifications, class enrichment and sensitivity to the lipid-annotation unit.

**Table S3a. Detected and qualifying $^{13}$C-labeled lipid annotations after host $^{13}$C-oleate feeding**

| Lipid class | Detected | Labeled | Labeled (%) |
|---|---|---|---|
| CL | 4 | 0 | 0 |
| Cer | 35 | 1 | 2.857 |
| CoQ | 2 | 0 | 0 |
| DG | 4 | 0 | 0 |
| DGCC | 10 | 1 | 10 |
| DGDG | 26 | 12 | 46.15 |
| DGGA | 1 | 0 | 0 |
| DLCL | 2 | 0 | 0 |
| FA | 27 | 4 | 14.81 |
| HBMP | 1 | 1 | 100 |
| HexCer | 1 | 0 | 0 |
| LPC | 21 | 3 | 14.29 |
| LPE | 29 | 4 | 13.79 |
| LPG | 1 | 0 | 0 |
| LPS | 3 | 0 | 0 |
| MGDG | 30 | 5 | 16.67 |
| MLCL | 1 | 0 | 0 |
| NAE | 7 | 0 | 0 |
| PC | 122 | 43 | 35.25 |
| PE | 84 | 20 | 23.81 |
| PEtOH | 1 | 0 | 0 |
| PG | 29 | 6 | 20.69 |
| PI | 34 | 5 | 14.71 |
| PS | 14 | 1 | 7.143 |
| SHexCer | 6 | 1 | 16.67 |
| SL | 9 | 4 | 44.44 |
| SM | 9 | 0 | 0 |
| SMGDG | 2 | 0 | 0 |
| SQDG | 28 | 1 | 3.571 |
| ST | 2 | 2 | 100 |
| TG | 217 | 74 | 34.1 |
| VAE | 10 | 1 | 10 |

**Table S3b. Effect of lipid-annotation unit on $^{13}$C-labeling counts and class enrichment**

| Annotation unit | Labeled | Detected | Enriched classes (BH-adjusted *P* < 0.05) |
|---|---|---|---|
| unique base/sum-composition annotation | 165 | 631 | Cer; TG |
| structural entry | 189 | 772 | Cer; PC; SQDG; TG |

**Table S3c. Fractional $^{13}C$ labeling by lipid class among qualifying annotations**

| Lipid class | Labeled | Mean labeling (%) | SEM (%) |
|---|---|---|---|
| Cer | 1 | 7.449 | 0 |
| DGCC | 1 | 11.99 | 0 |
| DGDG | 12 | 12.57 | 2.718 |
| FA | 4 | 7.699 | 0.9916 |
| HBMP | 1 | 11.12 | 0 |
| LPC | 3 | 34.14 | 12.05 |
| LPE | 4 | 31.93 | 7.176 |
| MGDG | 5 | 33.54 | 10.78 |
| PC | 43 | 26.76 | 4.406 |
| PE | 20 | 14.11 | 2.012 |
| PG | 6 | 15.98 | 5.054 |
| PI | 5 | 21.13 | 5.933 |
| PS | 1 | 6.597 | 0 |
| SHexCer | 1 | 29.6 | 0 |
| SL | 4 | 13.2 | 2.075 |
| SQDG | 1 | 6.227 | 0 |
| ST | 2 | 8.012 | 2.176 |
| TG | 74 | 27.38 | 2.674 |
| VAE | 1 | 9.262 | 0 |

**Table S3d. $NaH^{13}CO_3$-tracing feature classifications by biological compartment**

| Compartment | Source label | Named features | Statistically testable features | Qualifying labeled features | Nonqualifying features | Indeterminate features | % of testable features | % of named features |
|---|---|---|---|---|---|---|---|---|
| Algal metabolome | Algal metabolites: $NaH^{13}CO_3$ | 1127 | 1121 | 220 | 901 | 6 | 19.63 | 19.52 |
| Algal lipidome | Algal lipids: $NaH^{13}CO_3$ | 683 | 624 | 44 | 580 | 59 | 7.051 | 6.442 |
| Host cytosol metabolome | Host cytosol metabolites: $NaH^{13}CO_3$ | 1127 | 1121 | 52 | 1069 | 6 | 4.639 | 4.614 |
| Host LD lipidome | Host LD lipids: $NaH^{13}CO_3$ | 646 | 646 | 0 | 646 | 0 | 0 | 0 |

**Table S4. Algal TEM lipid-droplet measurements and lipidome composition statistics**

Supporting-data source: Supporting Data 3, A and C-D; total lipid values in Table S4b.

Cell-level lipid-droplet area and lipidome abundance, composition and annotation-level effects were compared between symbiotic and free-living algae. Oleic-acid treatment is stated explicitly where applicable. Relative-abundance tests for the ten classes displayed in Figure 4d and absolute class-summed abundance tests across all 24 detected classes are reported separately.

**Table S4a. Algal lipid-droplet area as a percentage of cell area in untreated and oleic-acid-treated free-living and endosymbiotic algae. Each analyzed cell was treated as an experimental unit.**

| Group | n | Mean LD area (% of cell area) | SD (%) | SEM (%) |
|---|---|---|---|---|
| Free-living | 30 | 0.6973 | 0.9205 | 0.1681 |
| Free-living + OA | 30 | 19.43 | 8.816 | 1.61 |
| Symbiotic | 30 | 3.385 | 1.837 | 0.3354 |
| Symbiotic + OA | 30 | 42.4 | 8.185 | 1.494 |

**Table S4b. Total algal lipid signal in symbiotic and free-living biological replicates**

Normalized total lipid signal for the eight algal biological replicates shown in Figure 4c.

| Algal state | Biological replicate | Total lipid signal (a.u.) |
|---|---|---|
| Symbiotic | SM_1 | 38378759839.3125 |
| Symbiotic | SM_2 | 35961023720.98947 |
| Symbiotic | SM_3 | 35132810387.11656 |
| Symbiotic | SM_4 | 36189754800.277336 |
| Free-living | FL_1 | 10936374432.210665 |
| Free-living | FL_2 | 9333111931.664627 |
| Free-living | FL_3 | 6928102268.18636 |
| Free-living | FL_4 | 8942538499.690033 |

**Table S4c. Numbers of the 633 unique algal lipid annotations detected across 24 lipid classes in the comparative symbiotic and free-living lipidomics dataset.**

| Lipid class | Detected annotations |
|---|---|
| TG | 229 |
| PE | 56 |
| PC | 52 |
| EtherPC | 38 |
| SL | 27 |
| MGDG | 25 |
| EtherTG | 24 |
| FA | 21 |
| EtherPE | 20 |
| DGDG | 19 |
| SQDG | 19 |
| PI | 18 |
| PG | 18 |
| LPE | 15 |
| EtherLPC | 12 |
| LPC | 12 |
| LPI | 9 |
| Cer_AP | 5 |
| PS | 4 |
| Cer_NP | 3 |
| CL | 3 |
| Cer_NDS | 2 |
| MLCL | 1 |
| LPG | 1 |

**Table S4d. Welch tests of relative abundance for the ten algal lipid classes displayed in Figure 4d. Percentages were calculated within each biological replicate, and *P* values were adjusted across the ten displayed classes using the Benjamini-Hochberg method.**

| Lipid class | n (symbiotic) | n (free-living) | *P* value | BH-adjusted *P* | Significance |
|---|---|---|---|---|---|
| DGDG | 4 | 4 | 0.01241 | 0.01379 | * |
| EtherLPC | 4 | 4 | $1.532 \times 10^{-5}$ | $5.175 \times 10^{-5}$ | **** |
| EtherPC | 4 | 4 | $1.552 \times 10^{-5}$ | $5.175 \times 10^{-5}$ | **** |
| FA | 4 | 4 | 0.3846 | 0.3846 | ns |
| MGDG | 4 | 4 | 0.002474 | 0.003534 | ** |
| PC | 4 | 4 | 0.01157 | 0.01379 | * |
| PE | 4 | 4 | $6.882 \times 10^{-4}$ | 0.001147 | ** |
| PG | 4 | 4 | $1.340 \times 10^{-4}$ | $2.680 \times 10^{-4}$ | *** |
| SQDG | 4 | 4 | $6.667 \times 10^{-5}$ | $1.667 \times 10^{-4}$ | *** |
| TG | 4 | 4 | $1.323 \times 10^{-6}$ | $1.323 \times 10^{-5}$ | **** |

**Table S4e. Algal lipid-class effects expressed as $\log_2$(symbiotic/free-living) from class-summed abundance. Welch tests were performed on $\log_2$(signal + 1), with Benjamini-Hochberg adjustment across all 24 detected classes.**

| Lipid class | $\log_2$FC (symbiotic/free-living) | *P* value | BH-adjusted *P* | Significance |
|---|---|---|---|---|
| Cer_AP | 0.8415 | 0.001577 | 0.002366 | ** |
| Cer_NDS | 4.077 | 0.009687 | 0.01057 | * |
| Cer_NP | 5.014 | $4.232 \times 10^{-5}$ | $1.263 \times 10^{-4}$ | *** |
| CL | 7.631 | $6.422 \times 10^{-5}$ | $1.541 \times 10^{-4}$ | *** |
| DGDG | 1.362 | 0.002791 | 0.003722 | ** |
| EtherLPC | 14.75 | $8.844 \times 10^{-8}$ | $7.075 \times 10^{-7}$ | **** |
| EtherPC | 7.8 | $3.192 \times 10^{-7}$ | $1.758 \times 10^{-6}$ | **** |
| EtherPE | 0.001902 | 0.8229 | 0.8229 | ns |
| EtherTG | 8.722 | $1.316 \times 10^{-5}$ | $4.512 \times 10^{-5}$ | **** |
| FA | 1.815 | $7.666 \times 10^{-5}$ | $1.673 \times 10^{-4}$ | *** |
| LPC | 2.08 | $4.736 \times 10^{-5}$ | $1.263 \times 10^{-4}$ | *** |
| LPE | 7.678 | $4.333 \times 10^{-11}$ | $1.040 \times 10^{-9}$ | **** |
| LPG | 2.858 | $4.728 \times 10^{-4}$ | $8.106 \times 10^{-4}$ | *** |
| LPI | 2.504 | 0.001298 | 0.002077 | ** |
| MGDG | 1.088 | 0.005674 | 0.006809 | ** |
| MLCL | 8.893 | $2.929 \times 10^{-9}$ | $3.515 \times 10^{-8}$ | **** |
| PC | 1.368 | $4.253 \times 10^{-4}$ | $7.851 \times 10^{-4}$ | *** |
| PE | -1.551 | 0.005998 | 0.006855 | ** |
| PG | 0.7055 | 0.002526 | 0.003566 | ** |
| PI | 1.881 | $6.037 \times 10^{-7}$ | $2.415 \times 10^{-6}$ | **** |
| PS | 0.4423 | 0.004068 | 0.005138 | ** |
| SL | 7.014 | $3.662 \times 10^{-7}$ | $1.758 \times 10^{-6}$ | **** |
| SQDG | 0.03168 | 0.8174 | 0.8229 | ns |
| TG | 5.315 | $1.910 \times 10^{-4}$ | $3.820 \times 10^{-4}$ | *** |

Significance codes reflect BH-adjusted *P* (q-value): ns, $P \geq 0.05$; *$P < 0.05$; **$P < 0.01$; ***$P < 0.001$; ****$P < 0.0001$.

**Table S5. Transcriptome-wide differential expression and lipid-pathway remodeling in symbiotic *Chlorella variabilis***

Supporting-data source: Supporting Data 4, A–G

Table S5 identifies the genes labeled in Supporting Figure S6 and summarizes the lipid-metabolic reactions displayed in Figure 5a, the direction of transcript-abundance differences in symbiotic relative to free-living algae, and the measured lipid-class endpoints. The reconstruction contains 93 displayed reaction-gene assignments representing 90 unique genes. Seventy-one expression-filtered reaction-gene assignments represented 69 unique genes; 34 had higher transcript abundance, 12 had lower transcript abundance and 23 did not meet both differential-expression criteria. A gene assigned to more than one reaction is counted once in each relevant reaction row.

**Table S5a. Functional identities and differential-expression results for the 16 genes labeled in Supporting Figure S6. Fold changes compare symbiotic with free-living algae. Higher and lower classifications require an absolute $\log_2$ fold change of at least 1 and a BH-adjusted P value below 0.05.**

| Figure label / transcript ID | Functional description | Reaction | $\log_2$ fold change | BH-adjusted *P* | Classification |
|---|---|---|---|---|---|
| ACCase CT-alpha g12395.t1 | Acetyl-CoA carboxylase carboxyltransferase alpha subunit, chloroplastic | Fatty-acid synthesis initiation | +4.85 | $3.88 \times 10^{-8}$ | Higher |
| KAS II g10049.t1 | 3-Oxoacyl-ACP synthase II, chloroplastic | Fatty-acid elongation | +4.51 | $4.35 \times 10^{-6}$ | Higher |
| Mitochondrial KAS g10101.t1 | 3-Oxoacyl-ACP synthase, mitochondrial | Fatty-acid elongation | +6.35 | $2.82 \times 10^{-9}$ | Higher |
| LACS4 g11457.t1 | Long-chain acyl-CoA synthetase 4 | Long-chain fatty-acid activation | +3.22 | $2.90 \times 10^{-7}$ | Higher |
| KCR g5406.t1 | Very-long-chain 3-oxoacyl-CoA reductase 1 | ER fatty-acid elongation | +5.91 | $2.46 \times 10^{-8}$ | Higher |
| Omega-3 desaturase g2967.t1 | sn-2 acyl-lipid omega-3 desaturase, chloroplastic | Fatty-acid desaturation | +6.21 | $3.69 \times 10^{-9}$ | Higher |
| GPAT/ATS1 g5496.t1 | Glycerol-3-phosphate acyltransferase ATS1, chloroplastic | Glycerol-3-P to LPA | +5.00 | $4.58 \times 10^{-9}$ | Higher |
| ATS2/LPAT g2461.t1 | 1-Acyl-sn-glycerol-3-phosphate acyltransferase ATS2 | LPA to PA | −1.72 | $5.75 \times 10^{-8}$ | Lower |
| PAP g6177.t1 | Phosphatidate phosphatase, LPP-alpha | PA to DAG | +3.64 | $1.28 \times 10^{-7}$ | Higher |
| DGAT2 g7979.t1 | Acyl-CoA:diacylglycerol acyltransferase 2 | DAG to TG | −3.42 | $3.78 \times 10^{-7}$ | Lower |
| CCT1/2 g11186.t1 | Choline-phosphate cytidylyltransferase 1/2 | PC biosynthesis | +0.60 | $1.20 \times 10^{-3}$ | Does not meet both criteria |
| PECT1 g12394.t1 | Ethanolamine-phosphate cytidylyltransferase | PE/PS biosynthesis | +2.04 | $1.27 \times 10^{-5}$ | Higher |
| PGPS1/2 g9095.t1 | CDP-diacylglycerol:glycerol-3-phosphate phosphatidyltransferase 1, chloroplastic/mitochondrial | PG biosynthesis | +0.96 | $1.58 \times 10^{-3}$ | Does not meet both criteria |
| DGD1/2 g2363.t1 | Digalactosyldiacylglycerol synthase 1/2, chloroplastic | DGDG biosynthesis | +0.10 | 0.526 | Does not meet both criteria |
| SQD2 g5055.t1 | UDP-sulfoquinovose:diacylglycerol sulfoquinovosyltransferase | SQDG biosynthesis | +1.17 | $4.95 \times 10^{-6}$ | Higher |
| ACX3 g8503.t1 | Acyl-CoA oxidase 3, peroxisomal | Beta-oxidation | −4.13 | $1.17 \times 10^{-7}$ | Lower |

**Table S5b. Reaction-level counts for the lipid-pathway reconstruction. Mapped counts are reaction-gene assignments rather than unique-gene counts because a gene may contribute to more than one reaction. Higher and lower categories require an absolute $\log_2$ fold change of at least 1 and a BH-adjusted *P* value below 0.05.**

| Reaction group | Mapped assignments | Expression-filtered assignments | Higher in symbiotic algae | Lower in symbiotic algae |
|---|---|---|---|---|
| PG synthesis | 6 | 5 | 0 | 0 |
| MGDG/DGDG synthesis | 2 | 2 | 0 | 0 |
| SQDG synthesis | 4 | 2 | 1 | 0 |
| GPAT | 3 | 3 | 1 | 0 |
| LPAT | 5 | 2 | 0 | 1 |
| PAP | 4 | 3 | 2 | 0 |
| DGAT | 9 | 5 | 2 | 3 |
| PC synthesis | 2 | 1 | 0 | 0 |
| PE/PS synthesis | 8 | 5 | 1 | 2 |
| Beta-oxidation | 13 | 9 | 3 | 3 |
| FA synthesis initiation | 7 | 5 | 4 | 0 |
| FA activation | 5 | 5 | 3 | 1 |
| FA elongation | 9 | 9 | 8 | 0 |
| ER elongation | 7 | 6 | 2 | 2 |
| Desaturation | 9 | 9 | 7 | 0 |

**Table S5c. Measured lipid-class abundance endpoints shown at the termini of the Figure 5a pathway reconstruction. Fold changes compare symbiotic with free-living algae. Welch tests used untransformed class-summed signals, with Benjamini-Hochberg adjustment across the 24 lipid classes.**

| Lipid class | Symbiotic mean class-summed signal | Free-living mean class-summed signal | $\log_2$ fold change (symbiotic/free-living) | Welch *P* | BH-adjusted *P* |
|---|---|---|---|---|---|
| CL | $1.624 \times 10^{6}$ | 8195 | 7.631 | $2.189 \times 10^{-4}$ | $3.753 \times 10^{-4}$ |
| Cer_AP | $2.776 \times 10^{7}$ | $1.549 \times 10^{7}$ | 0.8415 | $3.932 \times 10^{-4}$ | $6.291 \times 10^{-4}$ |
| Cer_NDS | $2.120 \times 10^{5}$ | $1.256 \times 10^{4}$ | 4.077 | $1.221 \times 10^{-7}$ | $1.465 \times 10^{-6}$ |
| Cer_NP | $1.034 \times 10^{7}$ | $3.200 \times 10^{5}$ | 5.014 | $6.075 \times 10^{-5}$ | $1.620 \times 10^{-4}$ |
| DGDG | $1.258 \times 10^{9}$ | $4.894 \times 10^{8}$ | 1.362 | 0.01977 | 0.02386 |
| EtherLPC | $5.569 \times 10^{9}$ | $2.021 \times 10^{5}$ | 14.75 | $2.162 \times 10^{-6}$ | $1.730 \times 10^{-5}$ |
| EtherPC | $3.665 \times 10^{9}$ | $1.645 \times 10^{7}$ | 7.8 | $9.712 \times 10^{-9}$ | $2.331 \times 10^{-7}$ |
| EtherPE | $1.970 \times 10^{7}$ | $1.967 \times 10^{7}$ | 0.001902 | 0.9942 | 0.9942 |
| EtherTG | $2.683 \times 10^{8}$ | $6.352 \times 10^{5}$ | 8.722 | $3.386 \times 10^{-6}$ | $2.032 \times 10^{-5}$ |
| FA | $8.905 \times 10^{7}$ | $2.531 \times 10^{7}$ | 1.815 | $8.503 \times 10^{-5}$ | $2.041 \times 10^{-4}$ |
| LPC | $6.390 \times 10^{7}$ | $1.512 \times 10^{7}$ | 2.08 | $5.766 \times 10^{-5}$ | $1.620 \times 10^{-4}$ |
| LPE | $1.348 \times 10^{8}$ | $6.581 \times 10^{5}$ | 7.678 | $5.338 \times 10^{-5}$ | $1.620 \times 10^{-4}$ |
| LPG | $7.062 \times 10^{5}$ | $9.741 \times 10^{4}$ | 2.858 | 0.007934 | 0.01058 |
| LPI | $6.367 \times 10^{4}$ | $1.123 \times 10^{4}$ | 2.504 | 0.04434 | 0.04838 |
| MGDG | $9.277 \times 10^{8}$ | $4.364 \times 10^{8}$ | 1.088 | 0.01988 | 0.02386 |
| MLCL | $3.333 \times 10^{5}$ | 700.1 | 8.895 | $1.781 \times 10^{-4}$ | $3.288 \times 10^{-4}$ |
| PC | $9.839 \times 10^{9}$ | $3.812 \times 10^{9}$ | 1.368 | $5.775 \times 10^{-6}$ | $2.772 \times 10^{-5}$ |
| PE | $1.177 \times 10^{9}$ | $3.448 \times 10^{9}$ | -1.551 | 0.02179 | 0.02491 |
| PG | $4.800 \times 10^{8}$ | $2.943 \times 10^{8}$ | 0.7055 | 0.002362 | 0.003334 |
| PI | $4.853 \times 10^{7}$ | $1.318 \times 10^{7}$ | 1.881 | $9.703 \times 10^{-5}$ | $2.117 \times 10^{-4}$ |
| PS | $8.619 \times 10^{6}$ | $6.343 \times 10^{6}$ | 0.4423 | 0.001571 | 0.002356 |
| SL | $1.079 \times 10^{8}$ | $8.345 \times 10^{5}$ | 7.014 | $1.301 \times 10^{-4}$ | $2.602 \times 10^{-4}$ |
| SQDG | $1.268 \times 10^{8}$ | $1.240 \times 10^{8}$ | 0.03168 | 0.8303 | 0.8664 |
| TG | $1.259 \times 10^{10}$ | $3.163 \times 10^{8}$ | 5.315 | $7.558 \times 10^{-6}$ | $3.023 \times 10^{-5}$ |

**Table S6. CD36-like candidate expression, localization predictions and inhibitor-associated phenotype**

Supporting-data source: Supporting Data 5, A; candidate expression values in Table S6a.

Biological-replicate TPM values summarize candidate expression in symbiotic and free-living algae. DeepLoc 2.1 predictions summarize the localization and membrane-associated features of the algal CD36-like candidate g8977.t1. Cell-level lipid-droplet area is also summarized for free-living algae exposed to oleic acid alone or together with SSO or BLT-1; each analyzed cell was treated as an experimental unit.

**Table S6a. CD36-like candidate transcript abundance in symbiotic and free-living algae**

Biological-replicate TPM values for candidate g8977.t1 shown in Figure 5b.

| Algal state | Biological replicate | TPM of g8977.t1 |
|---|---|---|
| Symbiotic | SM-1 | 4.79808 |
| Symbiotic | SM-2 | 5.925788 |
| Symbiotic | SM-3 | 4.431349 |
| Free-living | FL-1 | 0.539651 |
| Free-living | FL-2 | 0.725758 |
| Free-living | FL-3 | 0.607311 |

**Table S6b. DeepLoc 2.1 categorical localization prediction for CD36-like candidate g8977.t1**

| Protein ID | Predicted localization | Predicted sequence features | Predicted membrane class |
|---|---|---|---|
| g8977.t1 | Cell membrane | Signal peptide\|Transmembrane domain | Transmembrane |

**Table S6c. DeepLoc 2.1 compartment and feature probabilities for CD36-like candidate g8977.t1**

| Predicted compartment/feature | Probability |
|---|---|
| Transmembrane | 0.9175 |
| Cell membrane | 0.6216 |
| Lysosome/Vacuole | 0.4462 |
| Peripheral | 0.4424 |
| Endoplasmic reticulum | 0.3309 |
| Cytoplasm | 0.2967 |
| Golgi apparatus | 0.2914 |
| Soluble | 0.1743 |
| Lipid anchor | 0.1509 |
| Extracellular | 0.1206 |
| Nucleus | 0.0666 |
| Mitochondrion | 0.0564 |
| Plastid | 0.0246 |
| Peroxisome | 0.0039 |

**Table S6d. Lipid-droplet area as a percentage of cell area in free-living algae treated with oleic acid alone or together with SSO or BLT-1.**

| Treatment | Analyzed cells | Mean LD area (% of cell area) | SD | Median LD area (% of cell area) |
|---|---|---|---|---|
| Oleate-only control | 30 | 22.58 | 5.861 | 23.43 |
| SSO | 30 | 12.49 | 4.933 | 12.51 |
| BLT-1 | 30 | 12.53 | 6.229 | 12.64 |

| Comparison | Student's t *P* | BH-adjusted *P* |
|---|---|---|
| Oleate-only control vs SSO | $1.287 \times 10^{-9}$ | $2.574 \times 10^{-9}$ |
| Oleate-only control vs BLT-1 | $2.553 \times 10^{-8}$ | $2.553 \times 10^{-8}$ |

**Supporting Data**

**Supporting Data 1. Host lipidome replicate correlations, lipid-class composition and species-level measurements.**

**(A)** Host biological-replicate Pearson correlation matrix underlying Supporting Fig. S1a. **(B)** Replicate-level lipid-class percentages underlying Fig. 1c. **(C)** All 478 detected host lipid annotations and replicate abundances; statistical fields are reported for the 453 annotations eligible for the Fig. 1d fold-change analysis, and the remaining 25 are marked as not tested because one state had all-zero signal. The same abundance matrix underlies the Fig. 1a PCA.

**Supporting Data 2. Isotope-tracing measurements and analyses.**

**(A) Cell-level NanoSIMS $^{13}C/^{12}C$ measurements underlying Fig. 2d. (B)** Oleate-isotope labeling calls and isotopologue statistics underlying Fig. 3. **(C)** Unique-annotation and structural-entry class enrichment underlying Fig. 3b and Supporting Fig. S2b. **(D)** Bicarbonate-tracing feature classifications underlying Supporting Fig. S2e. The biological-replicate EA-IRMS $^{13}C$ atom percentages underlying Fig. 2e are in Supporting Table S2b.

**Supporting Data 3. Algal TEM and lipidome measurements and analyses.**

**(A)** Cell-level lipid-droplet area underlying Fig. 4b. **(B)** Algal biological-replicate Pearson correlation matrix underlying Supporting Fig. S3b. **(C)** Replicate-level lipid-class percentages underlying Fig. 4d. **(D)** Species-level abundances and statistics underlying Fig. 4e. **(E)** All 29 LipidOne indices underlying Supporting Fig. S4a-c. **(F)** Composition-normalized chain-unsaturation data underlying Supporting Fig. S4d-e. **(G)** Class-resolved chain-space effects underlying Supporting Fig. S4f-g. **(H)** Mean abundance, differential-abundance and host-fed oleate-isotope-overlap data underlying Fig. 4f. **(I)** Replicate abundance matrix for the 74 annotations supported by both isotope labeling and differential abundance underlying Supporting Fig. S5. The biological-replicate total lipid signals underlying Fig. 4c are in Supporting Table S4b.

**Supporting Data 4. Transcriptomic measurements, differential-expression results and functional annotations underlying Fig. 5a.**

**(A)** Transcript-level TPM values for three symbiotic and three free-living *Chlorella variabilis* biological replicates, together with condition-aware expression-filter status. **(B)** Complete limma results for the 8,479 transcripts that passed the expression filter, including mean expression, $\log_2$ fold change, moderated test statistic, raw *P* value, BH-adjusted *P* value and differential-expression classification. **(C)** Gene identifiers, functional descriptions, pathway assignments, expression values and differential-expression results for the 90 description-consistent lipid-pathway genes included in Fig. 5a; 69 passed the expression filter. **(D)** The 95 reaction-gene assignments considered for Fig. 5a, including two description-inconsistent automated assignments retained with an exclusion flag and reason. **(E)** Transcript abundance, $\log_2$ fold change, BH-adjusted *P* value and differential-expression classification for every assignment considered for Fig. 5a. **(F)** Gene descriptions and Mercator4/MapMan assignments for all predicted transcripts. **(G)** Gene-to-KEGG Orthology assignments obtained using GhostKOALA and DeepKOALA, including the assignment source.

**Supporting Data 5. Inhibitor-associated lipid-droplet phenotype.**

**(A)** Cell-level lipid-droplet measurements underlying Fig. 5e. Biological-replicate expression values for the CD36-like candidate shown in Fig. 5b are provided in Supporting Table S6a
